# Of Brobdingnag and Lilliput, or how island area may link body size, genome size and mutation rate

**DOI:** 10.64898/2026.05.11.724217

**Authors:** J. Rivas-Santisteban

**Affiliations:** Department of Biodiversity and Evolutionary Biology National Museum of Natural Sciences, CSIC, Madrid, Spain

**Keywords:** island rule, insular gigantism, insular dwarfism, genome size, drift-barrier hypothesis, Damuth’s law, Island Biogeography Theory, effective density

## Abstract

Insular gigantism and dwarfism are commonly framed by the island rule, whereby small species become larger and large species smaller. Yet why large species diminish, and why some taxa depart from this rule, remain unclear. Here I propose a neutral null model linking island area to body and genome evolution. Island Biogeography Theory predicts that smaller islands support lower census populations, while Damuth’s law predicts an inverse relationship between body mass and population density. If effective population size, *N_e_*, declines at a different rate from island area, effective density, *D_e_* = *N_e_/*area, will also change. Under the drift-barrier hypothesis, variation in *N_e_* may further influence genome size and mutation rate, *µ*. Island area may therefore jointly predict body mass, genome size and *µ*. Analyses of global and datasets and particular species are broadly consistent with this expectation, although the observed associations are generally modest.

## Introduction

Organisms do exhibit a colourful variety of physical properties. One property that has been recurrently studied is the size of the species. The species size and shape are governed by intrinsic causes (e.g. genotype, physiology) and extrinsic causes (e.g. nutrition). Thus, size variation (hereafter *κ*) is determined by multidimensional factors, making the falsification of its causes a daunting task.

Despite this limitation, consistent observations link *κ* in extant populations to potential causes. For example, insular speciation is often associated with changes in the body size of population X compared to that of X’s mainland relatives (Darwin, 1859*1964; Wallace, 1880*2018; Hinton, 1926; Foster, 1963, 1964; Keogh et al., 2005; McClain et al., 2006; Benton et al., 2010; Lomolino et al., 2013; Biddick et al., 2019; Sangster et al., 2025). These observations are commonly explained by the Foster’s or *island rule*, according to which some populations are more prone than others to undergo changes in body size during insular speciation. Such changes are termed “gigantism” when insular relatives are larger than their mainland counterparts (i.e. *κ >* 0), and “dwarfism” when they are found smaller (i.e. *κ <* 0) (Foster, 1963; Lomolino, 1985) Some observations appear to relate taxonomy; for example, while insectivores are not clearly affected by the island rule, rodents would be prone to gigantism (Foster, 1963, 1964). A more broad observation is that originally-small mainland species tend to undergo insular gigantism, while the reverse is true for originally-large mainland species –e.g. Fig. 1 from Lomolino (2005)– . However, the generality of the island rule seems to hang on the type of data examined (Meiri et al., 2006, 2008; Itescu et al., 2014).

**Figure 1:**
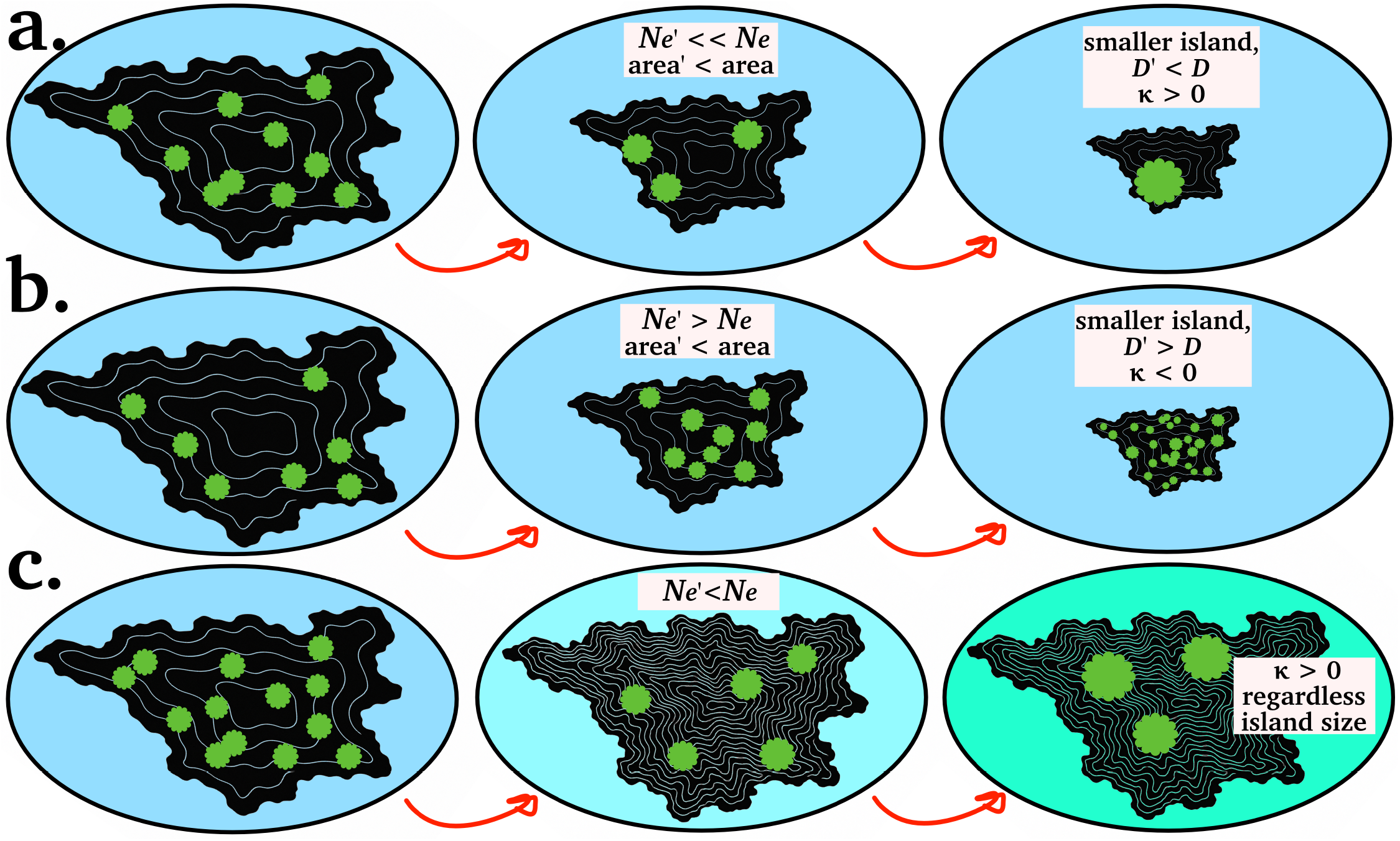
Predicted outcomes for *κ* following colonisation of progressively smaller islands under the core IBT model, Damuth’s law and nearly-neutral theory. a. Insular gigantism is stronger in small islands (*κ >* 0). This is a simple prediction, where *N_e_* of the species decreases more than island area, leading to increased body size as new effective density 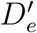 is lower. b. Insular dwarfism is stronger in smaller islands (*κ <* 0). This is often observed, although contrary to the naive IBT expectation where 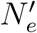 would decrease. Importantly, this scenario can still happen where 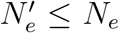 only if the smaller island provide Δ*area >* Δ*N_e_* . c. Other confounding factors can affect the prediction of body size using island size. In this example, insular stasis (*κ* ≈ 0) is expected since island size is almost equal. However, *N_e_* can be influenced by other factors, like unfavourable orography (number of isohypses in c.) or average temperature (background colour in c.), leading to *κ >* 0 in the example.

The evolutionary mechanisms underlying these observations remain unclear. First, under the core model of island biogeography theory (IBT) (MacArthur and Wilson, 1963; Warren et al., 2015), insular effects are expected to be stronger on smaller and more isolated islands. Island area influences extinction rates, whereas isolation constrains immigration. Together, these factors appear to predict the magnitude of the *island rule* (Damuth, 1993; Benítez-López et al., 2021; Vega-Rovira et al., 2025; Berg and Nietlisbach, 2025).

Second, a widely known explanation for the differential size of species is bound to the population census *N*, following Damuth’s law (Damuth, 1981). It is a simple explanation. The smaller the individuals of a population are, the higher number of individuals the population can maintain, and *vice versa*. It follows a negative power-law relationship,

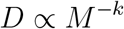

thus population density *D* declines with species’ body mass *M* at a rate *k. D* is simply 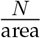.

Importantly, decreasing island area may reduce both census population size *N* and effective population size *N_e_* (Wright, 1931; Leroy et al., 2021). Let us consider an effective population density, defined as 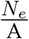, where *A* is a constraining island area (Fig. S1). Under this model, characteristic body size of a population, proxied by *M*, is expected to evolve only when decreases in *A* generate persistent change in population density, such that 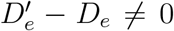 . This occurs when *N_e_* and *A* change at different rates, yielding a new effective population size 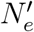 and consequently 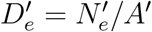 .The model is developed fully in Methods.

Thus, under this “effective density” model, three neutral scenarios are possible as island area decreases:

1. Both factors in the fraction 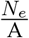 are equally diminished, rendering an equivalent effective density 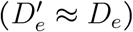 . Thus, neither gigantism or dwarfism is expected, but body size stasis (*κ ≈* 0).
2. Where 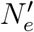 of the population decreased more profusely than the new island area. Here, the new population density is sparse 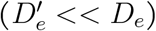, allowing for body size *M* increase (*κ >* 0).
3. If 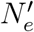 increases, remains equal, or decreases but the conquered island area is much more reduced than 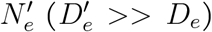, only populations with smaller body size can maintain the new higher density (*κ <* 0). Scenarios (ii) and (iii) may be sufficient to explain some cases of insular gigantism and dwarfism (Fig. 1a–b).

But these scenarios are not equally probable. If Damuth’s law and the IBT model are taken together, decreasing island area will more often constrain *N_e_* than expand it (Damuth, 1993). Ecological factors are also expected to interfere with this signal. Let me offer an example. The expansion of a mainland colonist on a small island might be limited if resources are permanently scarcer (Simberloff, 1976). *N_e_* would diminish 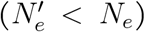 . But colonisation may increase 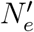 relative to mainland populations if reduced competition allows ecological release. Thus, these naive IBT expectations (smaller island = smaller *N_e_*) might leak some exceptions due to other factors (e.g. orography, competition (Pafilis et al., 2009); Fig. 1c)

### Box 1: Premises of the effective density model

**Island area limits how many individuals can exist (***N* )

Smaller islands have less habitats and fewer resources, thus a lower carrying capacity. On average, smaller island size limits more the census *N* . Following Damuth’s law, population density *D* is a proxy of the mass of an individual of that population, or *M* . Higher *D* predicts lower *M*.

**Census size (***N* **) shapes effective population size (***N_e_*)

When *N* collapses, *N_e_* typically collapses too (Kalinowski and Waples, 2002; Woolfit and Bromham, 2005; Leroy et al., 2021). On small islands, drift would become stronger on average if the population is effectively isolated (Fig. S1). Species adapted to larger islands with larger *N* keep higher *N_e_* and higher genetic diversity *θ*.

*N_e_* **informs** *µ* **and genome size**

These are predictions of the nearly-neutral theory and the drift-barrier hypothesis (Ohta, 1996; Sung et al., 2012). In small-*N_e_* populations, mildly deleterious mutations accumulate more easily, sometimes leading to genome expansion and increased *µ*. In large-*N_e_* populations, selection is more effective, *µ* can be lower or better optimised, and genome tends to be shorter. This assumption has recently been challenged in vertebrates (Marino et al., 2025; Weinstein and Roy, 2026) (see the “Limitations” section). Under this model, as long as *N_e_* variation recalls island area, genomic parameters are expected to covary with *N_e_* and island area.

**The combined effect provides a neutral null explanation for body size variation**

If viable body size strategies are derived from Damuth’s law, then reported insular *κ* could already be explained by the change in population density rather than by adaptive trends. This would be falsified by checking if island area covaries with Δ*N_e_*, thus variation in genome size and *µ*. In this study, I explore whether empirical data are compatible with such a model.

In this study, species body weights were compared across a global subset of 12,776 islands larger than 0.99 km^2^. Rather than asking whether insular populations are “giant” or “dwarf” relative to their closest mainland relatives, this study tests whether island area predicts average body size. If *κ* reflects a genuine response to insular constraints, this signal should emerge from comparisons across thousands of islands differing in area and their endemic biotas.

We can further test whether the *island rule* arises from changes in *D_e_* by interrogating genetic parameters (Box 1). Under nearly neutral theory and the drift-barrier hypothesis, lower *N_e_* may increase genome size and mutation rates (*µ*) (Kimura, 1991; Ohta, 1992, 1996; Lynch and Conery, 2003; Sung et al., 2012; Lynch et al., 2016). I therefore explore whether genome size covaries with island area, and whether *µ* covaries with island area or body size.

## Results and Discussion

### Island area predicts species richness

Briefly, for a subset the total Earth’s islands (12,776 with *>*0.99 km^2^, distribution of insular sizes available shown at Figure S2), a custom data-mining software was utilised to retrieve biological and geographic information linked to named islands from reliable global databases (see Methods). The smallest island in the dataset is Cherkgorsön, from the Alan islands (0.998 km^2^), and the largest is Greenland (2,108,459 km^2^). The number of named islands with biogeographic data was reduced but still important: 3,272 islands retained information for 120,907 species. As predicted by IBT, species richness increased with island area in this subset (Fig. S3).

### Island area predicts body mass

Studies of insular gigantism and dwarfism typically compare focal island species with their closest mainland relatives (Lomolino, 1985; Benítez-López et al., 2021). Here I instead ask how well island area predicts the mean body mass of all species recorded on each island (see Methods), thereby capturing the overall effect of island area, if any, across the largest possible dataset.

After retrieving the number of observations for each species across 3,272 named islands, the remaining variable was body mass. Holotype masses were compiled from Wikidata and the EOL TraitBank. Reliable estimates in grams were available for 1,942 species, representing only four animal classes: *Aves, Mammalia, Squamata* and *Insecta*.

The results show clear-but-modest trends (Fig. 2). Decreasing island size (geodesic area) partly predicted body weight in all four classes. *Aves* showed positive weight increases with decreasing island size (Fig. 2a & d), suggestive of positive *κ* in favour of the naive IBT expectation (Fig. 1a). On the contrary, *Squamata* showed a clear “counter-IBT” scenario; that is, the species found in smaller islands are increasingly small, on average (Fig. 2c). Mammals and insects showed only weak relationships, possibly because relatively few species were represented and the predominant taxa differed among islands, sometimes by orders of magnitude in body mass.

**Figure 2:**
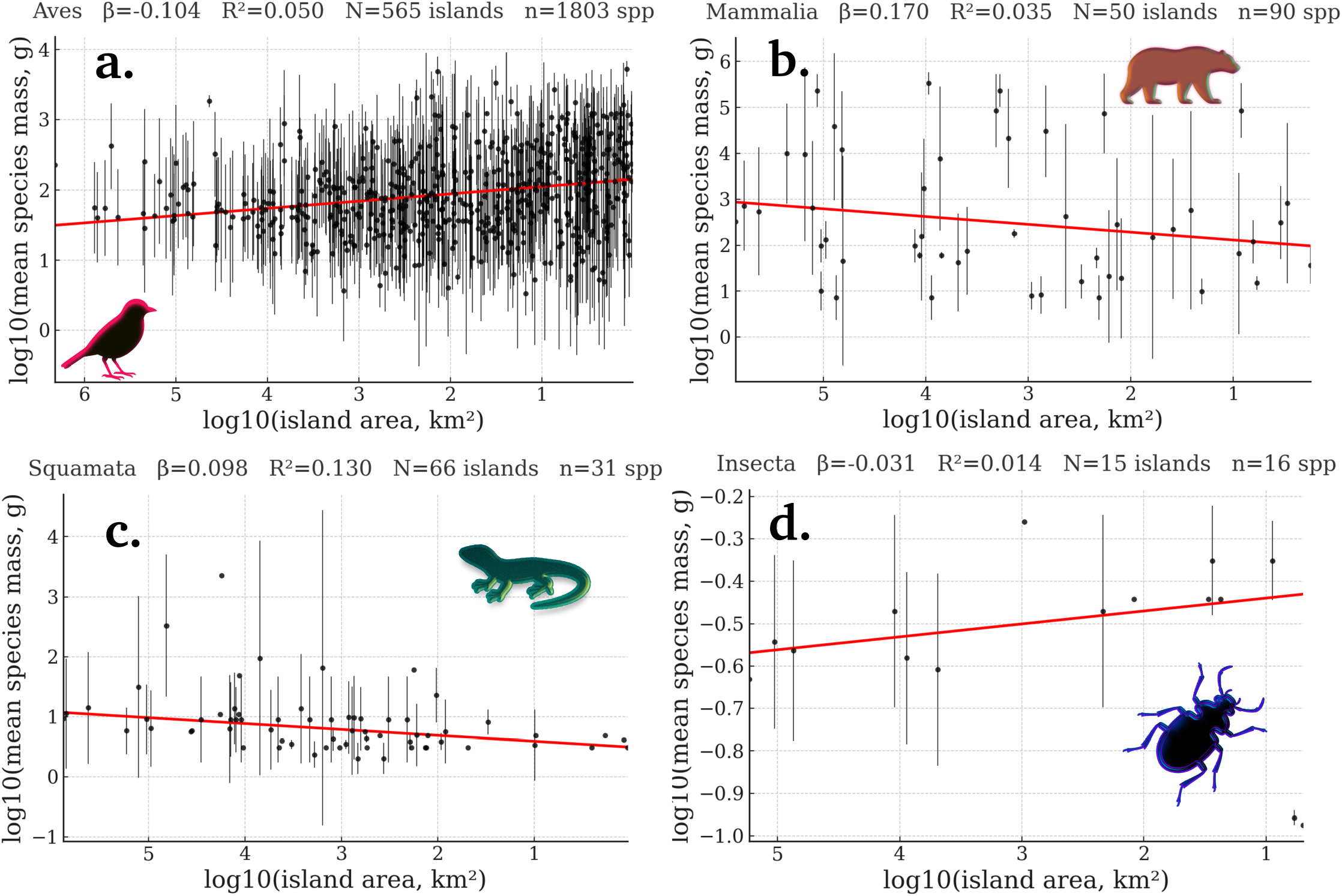
Island area predicts the mean body mass of species occurring on each island, although the direction of the relationship depends on taxonomy: (a) birds, (b) mammals, (c) squamates and (d) insects. Birds and insects follow the naive IBT expectation depicted in Figure 1a, whereby smaller islands support fewer but larger-bodied populations. Mammals and squamates show the opposite pattern, corresponding to the “counter-IBT” scenario in Figure 1b.

### Island area predicts body length

The relationships between island area and height, length and wingspan are shown. For the four classes examined above, these relationships were consistent with the body mass analyses, with the exception of *Insecta*, which now shows a “counter-IBT” scenario (Fig. S4). This discrepancy might be simply explained by a differential sample size: only weights of 16 species of insects over 15 islands were analysed, in contrast, lengths of 178 species over 191 islands are displayed.

Measurement data is available for other groups. It is shown that maximum height of flower plants decreases with decreasing island area, although sample size is small (Fig. 3a). In plants, the island rule is thought to strictly rely on ecology. For example, it was shown recently that wind-pollinated colonisers among archipelagoes in the SW Pacific tend to gigantism (compared to mainland), while animal-pollinated plants follow a strict island rule (Ciarle et al., 2025). I here contend that, although such ecological adaptations remain surely important, island area might exert an overlooked but independent effect over plant height evolution that can be neutrally explained by the effective density model (Fig. 1b).

**Figure 3:**
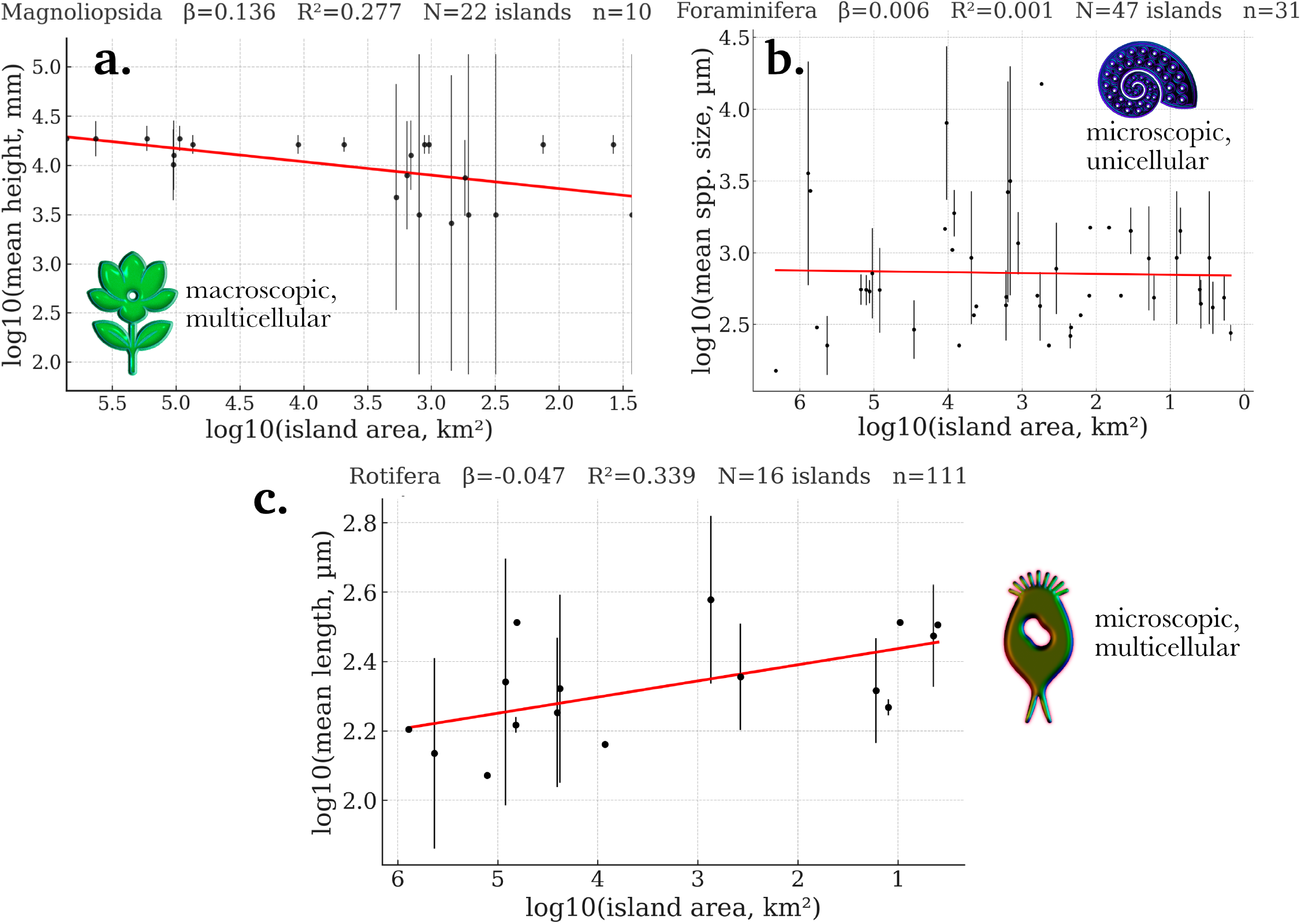
Decreasing island area predicts average species length regardless phylotypic size, in agreement with the interpretation that the slope polarity is dominated by Δ*N_e_* and Δ*A*. a: flower plants, b: forams, c: rotifers.

The other two groups examined are microorganisms. The first kind, *Foraminifera*, comprise a class of protists. Despite being unicelled, they exhibit a great range of sizes, from 0.02 mm to *>*20 cm in xenophyophores (Gooday et al., 2011). It is predicted that foram population densities *D_e_* are not affected by decreasing island size, so 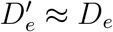, as these are ubiquitous within the coastal sediments (Murray, 2006; LeKieffre et al., 2017). Thus, body size stasis (*κ ≈* 0) is expected. The results indeed show no correlation between decreasing island area and average foram sizes for 31 species found across 47 islands (Fig. 3b). Thus, this negative result suggest that the effective density is a suitable model to predict whether insular gigantism or dwarfism is due in species where Δ*N_e_* ≠ Δ*area*, leading to differential 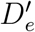.

Are there taxa in which the decline in *N_e_* is expected to outpace the decline in island area, thereby favouring island area-dependent gigantism? *Rotifera* can be a great case study. Most rotifers colonise freshwater habitats on islands, such as temporary ponds and lakes, and remain relatively isolated within them. However, because the number of freshwater habitats scales exponentially with island area and annual rainfall (De Manuel et al., 1992), rotifer *N_e_* may decline disproportionately on smaller islands, where such habitats become increasingly scarce. This would allow for gigantism. Results for 111 species across 16 islands support this prediction, revealing a clear positive *κ*: mean rotifer size increased significantly as island area decreased, with island area explaining 33.9% of the variation in mean size (Fig. 3c).

### Genome size and the *island rule*

It was shown how different observed proxies of body size might be compatible with a parameter *κ* responding to changes in effective density 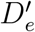 . However, if this model provides a useful null expectation, then (1) island area should partly predict genome size, and (2) mean body size across island biotas should correlate positively with mean genome size.

I provide a justification for both statements. In the effective density model developed here, changes in body size depend on *N_e_*, but also on the area effectively occupied by that idealised population. This only happens where changes in area are meaningful to *N_e_*, thus informing 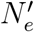 (Fig. S1). Under a nearly-neutral framework, such positive relationship between genome and body size is expected where genome expansion and contraction are primarily driven by shifts in *N_e_* (Ohta, 1992; Sung et al., 2012). That is to say, larger species will retain lower *N_e_* in a given area, potentially hinting genome expansion, while smaller species might undergo genome streamlining as their *N_e_* is higher. Exceptions to this expectation (Bobay and Ochman, 2018; Marino et al., 2025; Bastian et al., 2026; Weinstein and Roy, 2026) are discussed in “Limitations” section.

By constraining the *N_e_* of resident species to varying degrees, island area is expected to weakly predict the mean genome size of each island’s biota. This relationship should be weak because populations may be strongly, weakly or not at all constrained, depending on their spatial requirements (Fig. S1).

### Island area predicts genome size variation in endemic taxa

The following analyses use a substantially larger dataset because reliable body-mass and length data were less widely available. Genome sizes were retrieved through the NCBI Entrez API for 6,195 species, representing 5.12% of those listed.

Because some species occurred on multiple islands, we tested whether endemism influenced genome size variation (*κ*_*gen*_) (Fig. 4). The proportion of phyla showing “genome gigantism” declined as increasingly widespread species were included (Fig. 4a). When interrogating the behaviour of average *κ*_*gen*_ and its correlation level, it is shown that after the inclusion of species allocated to 4 islands or more, the signal of insularity on (*κ*_*gen*_) is almost null (Fig. 4b). Subsequent analyses were therefore restricted to endemic species, yielding 1,492 species across 225 islands.

**Figure 4:**
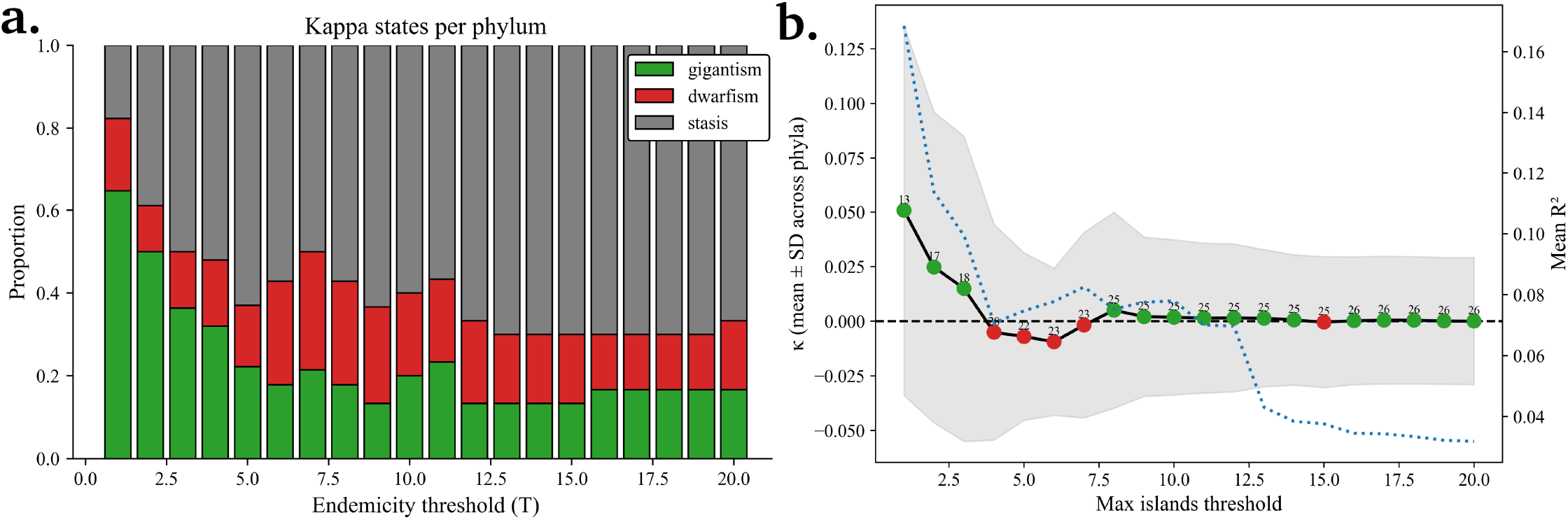
a. Proportion of phyla showcasing genomic “gigantism”, “dwarfism”, or “stasis” with decreasing island size across different endemicity thresholds. *T* = 1 includes only strict endemics, whereas *T* = 20 includes species occurring on up to 20 islands. b. Decay of *κ*_*gen*_ with decreasing endemicity. As species occurring in an increasing number of islands are included (numbers above the points indicate the number of represented phyla), *κ*_*gen*_ approaches an equilibrium, yielding null predictions. Note the concomitant decay in correlation at *T* = 4, 8, and 13 (blue dotted line).

Another consideration is that are different degrees of certainty for our data (e.g. number of species observations). Thus, regressions should be weighted by the total observations of every species within the islands. It is then assumed that if the same group of species of different genome size are found 10, 000 times in island X, and only 10 in island Y, island X’s area is 1,000-fold more informative for the genome size of the species set than that of the island Y.

The main result is shown in Figure 5, while complementary results are shown in Figs. S5 (by-island taxonomic composition), S6 (including std. deviations) and Table S1 (slope tests). The overall regression is represented by the red dotted line, and illustrates the extent to which a decrease in island area predicts variation in genome size. When considering all taxonomic levels, it has the form of a Simpson’s paradox (Hernán et al., 2011), as the three possible statistical classes described in Figure 1 were analysed together. When taxa are analysed separately, the regressions have substantially greater predictive power (Fig. 5). As anticipated by the effective density model, we see three scenarios: decreasing island area predicts increases in genome size for some taxa (*κ*_*gen*_ *>* 0; chordates, nonflying arthropods, and plants), a decrease for other (*κ*_*gen*_ *<* 0; flying arthropods), or genome size being unresponsive to island area (*κ*_*gen*_ *≈* 0; fungi). The below paragraphs briefly discuss how each outcome may be compatible with such a model.

**Figure 5:**
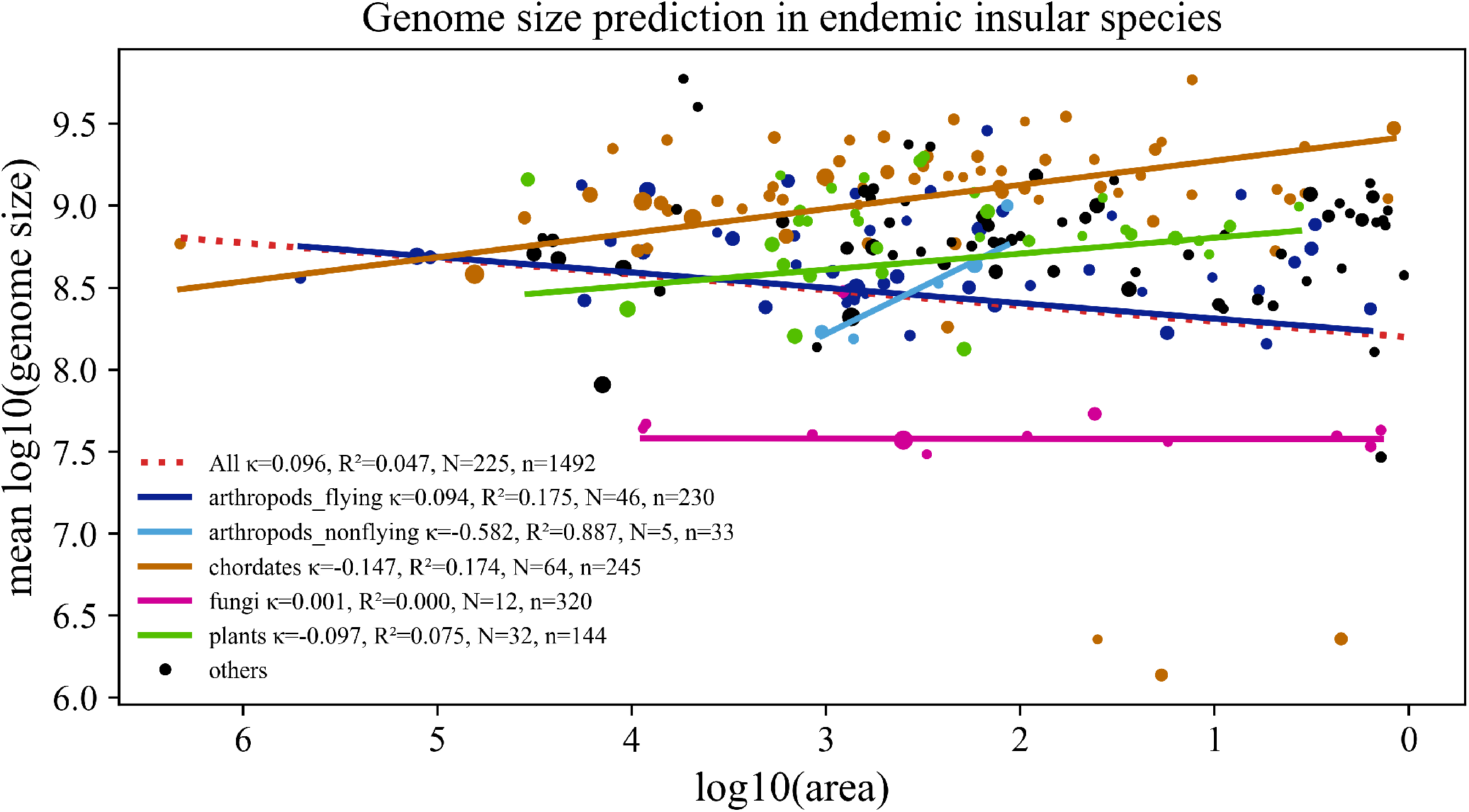
Mean genome size per island as predicted by island area, coloured by the predominant taxonomic group (Fig. S5). Each point represents an island, and point size reflects the number of species records contributing to its mean (standard deviations available in Fig. S6). *N* denotes the number of islands and *n* the number of endemic species with genome size data. As expected, island area was a substantially stronger predictor within taxa more prone to isolation, explaining up to 88.7% of the variance in non-flying arthropods, whereas no relationship was detected in fungi (*R*^2^ = 0). Pairwise comparisons of slopes are provided in Table S1.

Freshwater microfauna provided the clearest examples, as increases in body size should coincide only with genome gigantism. Rotifers and copepods showed larger genomes on smaller islands (*κ*_*gen*_ *>* 0, *R*^2^ *>* 0.4; Fig. S7), while rotifer body size increased too (*κ >* 0, *R*^2^ = 0.34; Fig. 3c).

Non-flying terrestrial arthropods should likewise experience disproportionate reductions in *N_e_*. Accordingly, springtails, millipedes and other non-flying arthropods had larger genomes on smaller islands (Figs. S8–S9), and so they reflect *κ*_*gen*_ *>* 0, *R*^2^ = 0.887; Fig. 5. Insects also tended to be heavier on smaller islands (Fig. 2d), although length showed the opposite trend (Fig. S4). Flying groups, particularly beetles, departed from this pattern, whereas *Lepidoptera* retained genomic gigantism (Fig. S8), perhaps reflecting their greater capacity for long-distance migration (Chapman et al., 2015).

Arachnids (*Arachnida*) exhibited, however, strong island area-dependent genomic stasis. This might be explained by the ability of some spiders to balloon (long-distance dispersal on silk threads carried by wind currents) (Rennie, 1831; Bishop, 1990; Szymkowiak et al., 2007). However, only 36.1% of analysed species could produce silk. Consistent with this interpretation, island area explained eightfold more variation in genome size among non-ballooning than ballooning arachnids (1.6% versus 0.2%; Fig. S9), although both effects were weak.

Mammals provide an interesting case. Carnivores and rodents show similarly predictable genomic gigantism as island area decreases, with area explaining 48.7 and 44.8% of the variance, respectively (Fig. S10), in agreement with recent observations (Berg and Nietlisbach, 2025). By contrast, chiropterans, despite overlapping rodents in body mass, show genomic dwarfism (30.3%; Fig. S10). This trend, shared with flying arthropods, might be reflecting that bats can easily migrate between islands, but not very distant islands (Fleming, 2019).

Plant responses may be particularly variable. Primary producers can colonise *de novo* islands without established communities. Thus, decreasing island area might, in some cases, increase 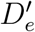 favouring smaller bodies (Fig. 1b). In these cases 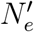 may increase, be indifferent, or decline. The tentative body size dwarfism (Fig. 3a) could be indeed compatible with area-dependent genomic gigantism, stasis, or dwarfism via 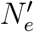, all of which occur among individual families (Fig. S11) irrespective of the overall genomic increase shown in Figure 5.

What about species whose population density is expected to remain decoupled from island area? *Scyphozoa* (true jellyfishes) are pelagic beings and should therefore face little limitation by island area. Indeed, they show an almost null relationship between island area and genome size (*R*^2^ = 1 × 10^−4^; Fig. S12). Other swimming classes were similarly unresponsive. Mail-cheeked fishes (*Scorpaeniformes*), whose habitats may depend more on sand or rock availability, show no correlation, perhaps because they can move freely between islands (*R*^2^ = 4 × 10^−4^; Fig. S12). Likewise, pedestrian classes such as *Malacostraca* were unresponsive to area (*R*^2^ = 6 × 10^−4^; Fig. S12).

Aves broadly conform to the model’s expectation, as area-dependent mass gigantism (Fig. 2a) only predicts genome gigantism, and evidence of genome gigantism is observed overall (Fig. S13). Thus, this preliminary, broad-brush analysis of the empirical data suggests that the observations are compatible with, or partially explained by a neutral null.

### Genome size of insular species is predicted by body mass

Another key prediction from combining Damuth’s law with the drift-barrier hypothesis is that body mass should correlate with genome size and mutation rate (Majic et al., 2025), where mass *M* increases are likely due to a diminution in species effective density 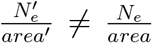 . Under the premises that larger organisms occur at lower population densities and that lower *N_e_* permits genome expansion, body and genome size should covary positively as area decreases. There are solely two exceptions: (1) where body size stasis occurs 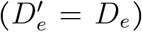 but *N_e_* diminish at the same rate than area, thus the expectancy is genome gigantism and higher mutation rates; or (2) body size dwarfism occurs due to 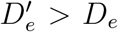 where 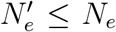, thus bounded to genome stasis (if 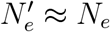 ), or to genome gigantism (if 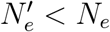 ).

The average weight in grams of every species found within each island was used as a predictor of the average genome size of the species observed in an island (Fig. 5a including widespread species, Fig. 5b limited to endemic species). As expected, a positive relationship is found. The fact that prediction is stronger when avoiding widespread species (*R*^2^ = 0.108) indicates that, as expected by such a model, by-island mean weight measures pertaining to nonendemics is less informative of by-island mean genome sizes than weight measures only pertaining to endemics.

### Evidence that area-driven Δ*D*_e_Δ*N*_e_ partly predicts *κ* and *κ*_gen_

For the drift-barrier hypothesis to be correct, it is expected that species with lower *N_e_* exhibit higher mutation rates *µ* (Lynch and Conery, 2003; Sung et al., 2012). As slightly deleterious mutations will accumulate “at random”, they will eventually damage the machinery that enables high-fidelity copying. This allegedly causes a feedback loop, allowing for more errors per site per generation, contributing to higher *µ*. Likewise, low *N_e_* naively predicts genome expansion (Lynch and Conery, 2003; Bobay and Ochman, 2017), although it is a complex relationship (Marino et al., 2025) (see Limitations).

In summary, provided that small islands constrain *N_e_* more than bigger islands on average, drift-barrier effects would tend to be stronger in the former. Changes in genomic features via drift-barrier effects will be linked to changes in body size via Damuth’s law where 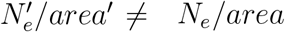.

### Case study 1: Darwin’s finches *N_e_* is dependent on island area

To study whether island size affects *N_e_*, we can make use of reliable data on the polymorphism of wild populations (derived from SNPs) of Darwin’s finches in the Galapagos archipelago (Lamichhaney et al., 2015). Correcting the SNPs by segregation sites (affected by sampling effort), Watterson’s *θ* can be extrapolated assuming a constant mutation rate from the ancestor, the zebra finch (*Taeniopygia guttata*’s *µ* = 2.04 × 10^−9^ per site per year) (Lamichhaney et al., 2015). Once *θ* is obtained, *N_e_* is deduced for each finch population on different islands. While it was previously recognized that *N_e_* depends on island size (Lamichhaney et al., 2015; Leroy et al., 2021), here I analyse whether these differences are common to different species and whether they can be predicted from differences in 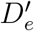 and/or area.

The effective density model predicts that the *N_e_* of Galapagos finches should be smaller on smaller islands. This is found only if Genovesa island is excluded (Fig. S14; Table 1). Genovesa can be considered an outlier for its very low immigration rates (Farrington and Petren, 2011), and selection to dry conditions (Grant, 1985), both factors leading to a lower variability. These factors may influence *N_e_* estimates (Fig 1c). Excluding Genovesa, these data are in agreement with our result of *κ >* 0 in *Aves*, supported by larger body size (Fig. 2a) and body length (Fig. S3a) with decreasing area. In addition, this interpretation is supported by the genome-enlargement *κ*_*gen*_ *>* 0 trend envisaged for *Anseriformes* (Fig. S13). If we include Genovesa, and data for every finch species, the *N_e_*-decreasing trend is still found (−3.15 + *N_e_*× km^−2^; Fig. S15). These results are in agreement with Johnson and Seger (2001); Woolfit and Bromham (2005), but are inconsistent with the faster evolutionary rates observed in some bird populations inhabiting larger landmasses (Wright et al., 2009).

**Table 1.** Exact permutation tests (two-sided) for Pearson correlations between raw variables in *G. difficilis*, excluding Genovesa (considered an outlier). For each predictor–response pair, r is the Pearson correlation coefficient computed on untransformed data, and *p*_perm_ is the exact permutation *p*-value based on all 5! = 120 permutations under the null hypothesis of no association. Area and *N_e_* are highly correlated. Data from (Lamichhaney et al., 2015).

| Test | Predictor | Response | $r$ (Pearson) | $p_{\text{perm}}$ |
| --- | --- | --- | --- | --- |
| 1 | Area | $N_e$ | 0.969 | 0.050 |
| 2 | $N_e$ | SNP | -0.193 | 0.783 |
| 3 | Area | SNP | 0.046 | 0.942 |

### Case study 2: *Chelonoidis* spp. mutation rates depend on body mass and island area

In the previous section it was shown that *N_e_* is prone to diminish with island area, an essential relationship for the model here proposed, although Δ*N_e_* is not deduced solely from Δ*area* as there could be confounding factors and gene flow between close islands.

However, the calculation of the long-term *N_e_* assumed a constant mutation rate *µ* = 2.04 × 10^−9^ as per Lamichhaney et al. (2015). We can now ask whether *µ* will collaterally change with Δ*N_e_*. As said, mutation rates *µ* will on average increase if 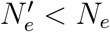, regardless of effective density changes.

Let us nonetheless study a clear case of 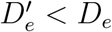 with decreasing island area, where the only prediction possible an increase in body size. In this scenario, the null is to consistently observe larger mutation rates *µ* at equilibrium, because *N_e_* must decrease more than area.

Preliminary support can be obtained by comparing neutral mutation rates among closely related species differing in body size. If *µ* correlates positively with body size and negatively with island area, drift-barrier effects on *µ* should be predictable from island area. An ideal species set should have reference genomes available, be restricted to oceanic islands isolated from continental landmasses, and exhibit lower migration rates than finches, allowing island area to constrain the effective area (Fig. S1).

Giant tortoises of the genus *Chelonoidis* provide a suitable system because their phylogeny and divergence times are well resolved (Jensen et al., 2021, 2022). *C. abingdonii* (Pinta, 60 km^2^), *C. phantasticus* (Fernandina, 642 km^2^), and the continental *C. denticulata* (Fig. S16 a–b) were selected. Although *C. niger* or *C. chilensis* would be preferable continental comparators, suitable genomes were available only for *C. denticulata*. The largest species, *C. abingdonii*, inhabited the smallest island, and its genome was sequenced from the last known individual, “Lonesome George” (Edwards et al., 2013).

Assuming *T* = 1.5 Myr (Poulakakis et al., 2020; Jensen et al., 2021), median branch-specific *d*_*S*_ values were used as coarse proxies of synonymous mutation rates as *µ ≈ d*_*S*_*/T* (Fig. S16 c–d). To exclude poorly resolved estimates and extreme outliers *d*_*S*_ range is restricted to 10^−5^–10^−2^. *µ* was estimated as 1.7 ×10^−9^, 8.9 ×10^−10^, and 7.7 ×10^−11^ substitutions per site per year for *C. abingdonii, C. phantasticus*, and *C. denticulata*, respectively. Thus, *C. abingdonii* evolved approximately 22-fold faster than *C. denticulata* and 2-fold faster than *C. phantasticus*. Using unfiltered median *d*_*S*_ values yielded a smaller 6-fold difference between *C. abingdonii* and *C. denticulata*.

Using body masses of 87.5 kg for *C. abingdonii*, 46.86 kg for *C. phantasticus* and 35 kg for *C. denticulata*, the corresponding lower-bound mutation rates are *µ/M* = 1.94 × 10^−11^, 1.9 × 10^−11^ and 2.2 × 10^−12^ mutations per site per year per kilogram, respectively (Fig. S16 e). Body-mass normalisation therefore makes *C. abingdonii* and *C. phantasticus* (insular species) almost indistinguishable in terms of synonymous mutation rate per kilogram (just 1.02-fold of difference), while both remain about 8.8-fold higher than *C. denticulata*, the closest continental ancestor available. These results are compatible with the effective density model.

### Further considerations

#### Density *D* vs. effective density *D_e_*

Damuth’s law describes an allometric relationship between population density and body mass, typically expressed as a census density *D* measured at a given point in time. This density *D* is *observational*, because it can be directly estimated from any extant population of individuals. In contrast, the effective density *D_e_* is a *theoretical* optimum. It is defined from the effective population size *N_e_* and the area occupied by it (effective area *A*; Fig. S1). *N_e_* is the number of individuals from an idealized population, and its numeric value is usually lower than the census *N* . However, it is relevant for evolution, because both long-term neutral genetic patterns and the extent of purifying selection are determined by *N_e_*, and thus better reflected by *D_e_*, rather than by transient fluctuations in census *N* or density *D*. Available area is known to influence both quantities, *N_e_* and *N* (Kalinowski and Waples, 2002; Leroy et al., 2021). Consequently, if body size variation is expected to arise in response to changes in population density within two populations belonging to the same species, such evolutionary responses should be more closely associated with *D_e_* than with the instantaneous census density *D*, because the latter can only account for variation between already-extant species. Slow changes in genome size and mutation rate through time will therefore be better captured by changes in *D_e_*. Instead, *D* cannot inform *κ*_*gen*_ because the latter is not an instantaneous quality. In this work, *D_e_* is approximated naively as the exact number of individuals per km^2^ required to achieve the species’ estimated *N_e_* on a given island.

The latter assumes that the original effective area was larger than the area of the colonised island (Fig. S1), which may explain why some island populations retain similar genetic diversity and estimated *N_e_* when compared to those of mainland (James et al., 2016).

In other words, *D_e_* is the equivalent density of breeding individuals per km^2^ in an ideal population – panmictic, demographically stable, without reproductive skew, and all the other assumptions for Fisher-Wright populations (Charlesworth, 2009)– that would experience the same genetic drift as the real population on that island.

### Could island area influence cell size?

The results herein suggest that microscopic organisms can likewise undergo area-dependent gigantism and dwarfism, provided Δ*D_e_*. One disregarded question is: could the island size be influencing cell size variation among unicells?

As population sizes of unicellular organisms are often dauntingly large when compared to those of multicellulars, and their densities are much higher as these larger populations can be confined within minuscule areas, area-dependent gigantism/dwarfism within those may seem, *a priori*, trivial questions to ask for. This might be the case because, additionally, many unicelled species are ubiquitous. This anticipation is supported by the no-relationship of foram body size with decreasing island area (Fig. 3b).

Indeed, we expect 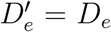 despite decreasing area, because island area greatly exceeds the effective area of single-celled organisms (Fig. S1). For example, consider a mammal effective population of *N_e_* = 10^4^. It will occupy 100 km^2^ of the island’s available surface if *D_e_* = 1, 000 individuals / km^2^. If we work out the maths for cyanobacteria, even with *N_e_* = 10^9^ and a density of 5 × 10^18^ cells / km^2^, the effectively occupied area, assuming no superposition, would be of just 0.2 m^2^. Thus cell size should be unresponsive to island area.

However, it is sensible to think of two special cases in which *N_e_* of unicells can be constrained by the size of the island: (1) unicells that are geographically isolated into highly specific areas, which abundance and extension exponentially depends on island area (e.g. volcanic alkaline lakes), (2) endosymbionts who are limited by the area of the island as is their host (Bobay and Ochman, 2017).

### Limitations

An important limitation is that neither remoteness nor distance between islands were considered, despite being fundamental to insular ecosystems (Warren et al., 2015), and often related to the *island rule* (Vega-Rovira et al., 2025). Given the heterogeneous sample sizes across taxa, these analyses should be interpreted primarily as exploratory. In addition, it does not account for the intricacies of radiation processes (Gillespie and Whittaker, 2025). Yet, despite being such a naive model, it allows for (weak) prediction (Figs. 2—6).

**Figure 6:**
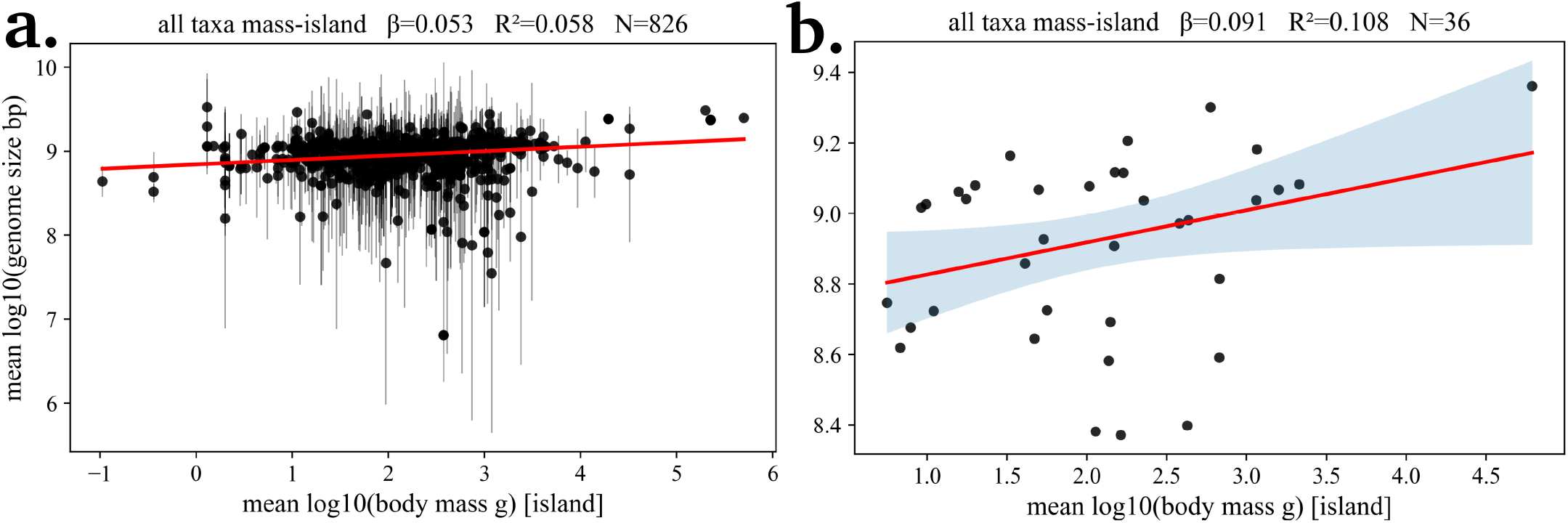
By-island average species’ genome size prediction by by-island average species’ weight. a. Widespread species allowed (species data for 826 islands). b. Only unique-island species allowed (species data for 36 islands). In both cases, prediction is positive. However, variance explanation achieved 10.8% using data for species found in only one island. Widespread species seem to add noise to the genome size prediction, as expected since their populations are not delimited by just one island.

Insular typology was neither considered, even if it is known to be an important driver of evolution. Wallace extensively discussed the important distinction made originally by Darwin on the origin of islands (Wallace, 1880*2018, 1887), which could be either “oceanic” (*de novo* island) or “continental” (*fragment* island) (Warren et al., 2015). This distinction is key, because oceanic islands emerge largely uncolonised, whereas continental islands inherit established biotas. However, it is less consequential here because if smaller islands exert stronger effects on *N_e_* this signal should remain detectable in the average traits of their inhabitants.

Although the nearly-neutral framework considers that genome expansion occurs where 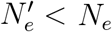, not every observation fits this prediction (Whitney et al., 2010; Ai et al., 2012; Zhang et al., 2024; Marino et al., 2025; Bastian et al., 2026). However, some of the latter exceptions were anticipated (Bobay and Ochman, 2017, 2018). Supplementary analyses of more global datasets further show that genome size and mutation rates strongly correlates with both *N_e_* and body mass, and that *N_e_* is dependent on mass (Fig. S17, data from Gibson et al. (2018); Bergeron et al. (2023); Wang and Obbard (2023); Beichman et al. (2024)).

Antropogenic disturbance of species-area relationship was not considered (Jardim de Queiroz et al., 2025). Hydrological parameters (e.g. rainfalls) were not considered, even if they influence the species-area relationship (Steibl et al., 2025). Generation overlap, temporal population structure, and skewed sex ratios may weaken the link between island area, *N* and *N_e_*. Incorporating these factors should improve lineage-specific predictions and might provide a useful direction for future work.

The synonymous rate estimates do not account for ancestral polymorphism contributing to divergence. Although recent inbreeding in *C. abingdonii* may have reduced contemporary heterozygosity, *d*_*S*_ primarily reflects long-term synonymous substitution accumulation (Bastian et al., 2026).

### Final notes

Here I propose the effective density model as a null expectation for patterns of variation during insular colonisation, linking island area, body size variation (*κ*) and genome length variation (*κ*_*gen*_). If germline mutation rates differ from the null expectation, selection becomes a more appropriate model to explain *κ* instead. Analyses of a global dataset, together with preliminary analyses of Galápagos tortoises and finches, can be broadly consistent with a neutral expectation, although the observed associations are generally weak.

A central assumption of the effective-density model is that effective population size *N_e_* varies with island area and, in turn, influences genome size and *µ*. Although these relationships are partly shaped by lineage-specific covariates (Marino et al., 2025; Weinstein and Roy, 2026), global datasets show that *N_e_* and body size explain significant fractions of variation in mutation rates and genome size (Fig. S17), consistent with expectations under changes in effective density.

## Methods

### Island and species data sets

Island polygons and areas were obtained from the UNEP–WCMC Global Islands database (ArcGIS FeatureServer, layer 2) and restricted to named islands *≥* 0.99, km^2^. Species inventories were retrieved from GBIF using polygon-constrained queries, whereas bodysize traits (mass, length, SVL, wingspan, etc.) were obtained from EOL TraitBank and, when unavailable, Wikidata. Of 12,776 islands retained, 3,272 contained biodiversity records, encompassing 120,907 species used in subsequent analyses.

### Island filtering and body size aggregation

Each occurrence was assigned to a taxonomic class, and body mass (g) was retrieved from the above databases. Mass was log_10_-transformed, so means represent geometric mean body mass. Data were aggregated by island and class, recording mean mass, its standard deviation, and species richness. Mean mass was then plotted against island area, with within-island standard deviations as error bars and the x-axis reversed from large to small islands.

For each taxon, we fitted a Huber-weighted robust linear model (RLM with HuberT),

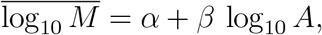

We report the robust slope *β*, ordinary least-squares *R*^2^, number of islands *N*, and number of species *n*. Because the x-axis is reversed for presentation, the visual direction of the trend is opposite to the sign of (*β*).

### Genome size retrieval

Genome size was retrieved (total assembled nuclear DNA length, in base pairs) for each species by programmatically querying NCBI genome assemblies. For every taxon in our island dataset, it was first resolved a valid binomial name (e.g. *Genus species*) and queried the NCBI Assembly database via Entrez to obtain up to 20 candidate genome assemblies. RefSeq assemblies (accessions starting with GCF ) were preferentially selected. If none were available, GenBank assemblies (GCA ) were selected. For each selected assembly, its NCBI FTP path was reconstructed, and downloaded the corresponding *accession* assembly stats.txt file, which is generated by NCBI for each assembly and reports the “total length” field, i.e. the total assembled sequence length in base pairs. This “total length” value was used as the genome size for that species.

### Whole-genome resequencing and consensus genomes

Paired-end whole-genome resequencing data from two *C. phantasticus* individuals from Fernandina Island (BioSamples SAMN 24674816 and SAMN 24674817) were retrieved from the SRA. Reads were quality-filtered and adapter-trimmed with fastp v1.0.1 (Chen et al., 2018), then aligned with BWA-MEM v0.7.19-r1273 to the *C. abingdonii* reference genome CheloAbing 2.0 (GCF 003597395.2; “Lonesome George”). BAM files were sorted and indexed with samtools v0.1.19 (Danecek et al., 2021), and mean coverage was estimated with “samtools depth”. Subsequent analyses used a *∼* 5× coverage library (SRR17619844), whereas the second served only to confirm segregating variants. Variants were called with bcftools v1.22 (Danecek et al., 2021) using “bcftools mpileup” and “bcftools call”, retaining sites with QUAL *≥* 30 and depth *≥* 4. Pseudohaploid consensus genomes were reconstructed with “bcftools consensus”. For *C. denticulata*, coding and protein sequences were extracted from assembly GCA 046118415.1 and its NCBI GFF3 annotation using gffread. Orthologues and the species tree were then inferred from the three species with OrthoFinder v3.0 (Emms and Kelly, 2019).

Lineage-specific synonymous substitution rates were estimated as median(*d*_*S*_)*/T*, using terminal branch *d*_*S*_ values inferred with “codeml” (PAML v4.10) (Yang, 2007). For some analyses, estimates were restricted to 10^−5^ *≤ d*_*S*_ *≤* 0.01, excluding near-zero values or highly divergent genes prone to alignment artefacts. T=1.5 Myr was assumed (Jensen et al., 2021). These values provide conservative proxies for lineage-specific mutation rates.

Body mass for each *Chelonoidis* species was approximated using the highest published estimate of mean adult male body mass.

### The effective density *D_e_* model

We define the effective density of breeding individuals as

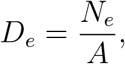

where *N_e_* is the long-term effective population size and *A* is the area occupied by *N_e_*. Naively, let us consider that it is larger than the area of the island, so the maximum value that an isolated species can effectively occupy is limited by the area of the island (Fig. S1). After a change in island size, we have

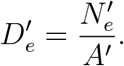

Combining these expressions yields

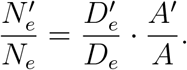

For shrinking islands (*A*^*′*^ *< A*), an increase in effective density 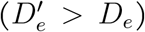 does not uniquely determine the direction of change in *N_e_*. Specifically, 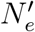 will decrease, remain unchanged, or increase if

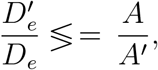

respectively. Thus, even if body size *κ* tends to decrease when *D_e_* increases, consistent with Damuth-like scaling of density with body mass, the long-term effective size *N_e_* may decline, remain stable, or increase, depending on the balance between changes in density and area.

Because density–body mass relationships and many life-history/allometric relationships are well approximated by power laws, we work on the log scale. Following Damuth’s law, we assume that long-term equilibrium body mass *M* obeys

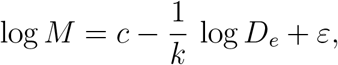

where *k >* 0 is the allometric exponent and *ε* collects other sources of variation. Therefore, the expected change in log body mass between two islands (or two demographic regimes) is

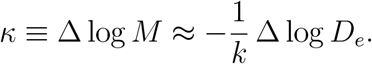

Thus, increases in effective density (Δ log *D_e_* > 0) tend to cause body dwarfism (*κ <* 0), whereas decreases in effective density (Δ log *D_e_* < 0) tend to cause body gigantism (*κ >* 0).

But, under which scenarios is the sign of *κ* the same that *κ*_*gen*_? Following the drift-barrier hypothesis, the long-term genome size *G* depends, on average, on *N_e_*. As a simple phenomenological approximation, we write

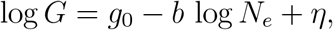

with *β >* 0 for lineages where lower *N_e_* allows larger genomes, and *η* capturing additional noise and lineage-specific effects. Consequently, the expected change in log genome size is

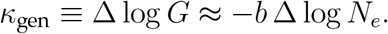

Thus, body size *κ* and genome size *κ*_*gen*_ will change in the same direction only if

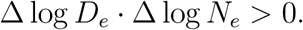

Using

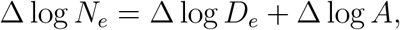

this condition can be written as

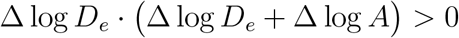

Biologically this implies that *κ* and *κ*_gen_ will retain the same polarity (e.g. species suffering of “gigantism” will suffer of genome expansion) whenever the change in effective density *D_e_* and the change in effective population size *N_e_* have the same sign, assuming that the new colonised island can be either bigger or smaller. Here it is assumed that new colonised island *A*^*′*^ was always of inferior size for narrative purposes.

### Map retrieval

Map data from [OpenStreetMap](https://www.openstreetmap.org/copyright), available under the Open Database License (ODbL).

