## Supplementary Information for "Of Brobdingnag and Lilliput, or how island area may link body size, genome size and mutation rate"

---

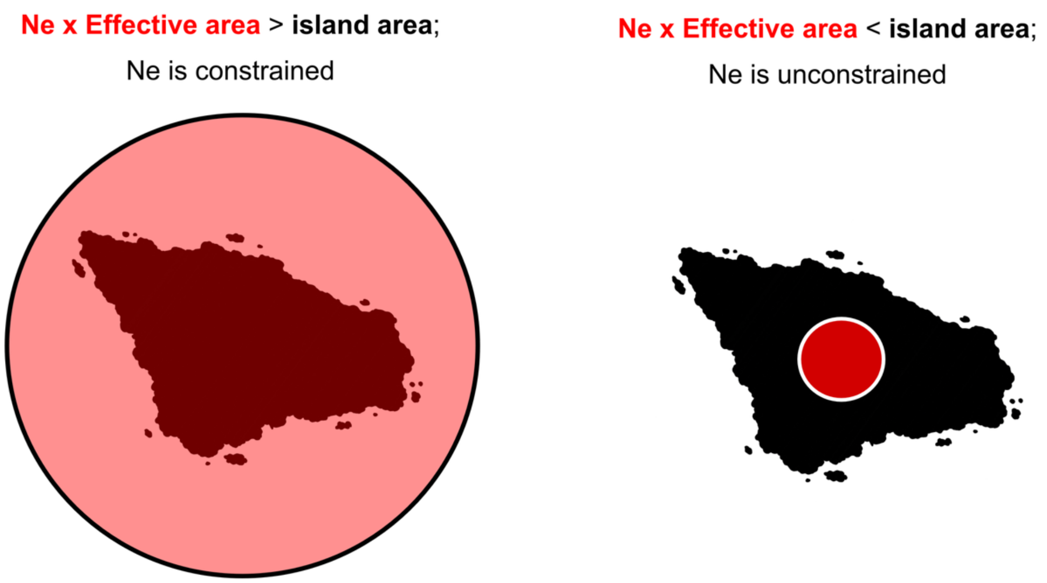


Figure S1. Concept of effective area, defined as the product of Ne and the area required per individual. Left: the species is spatially constrained by island area. Right: the species is not constrained by island area and is therefore unaffected by the effective-density model unless Ne or individual area requirements change.

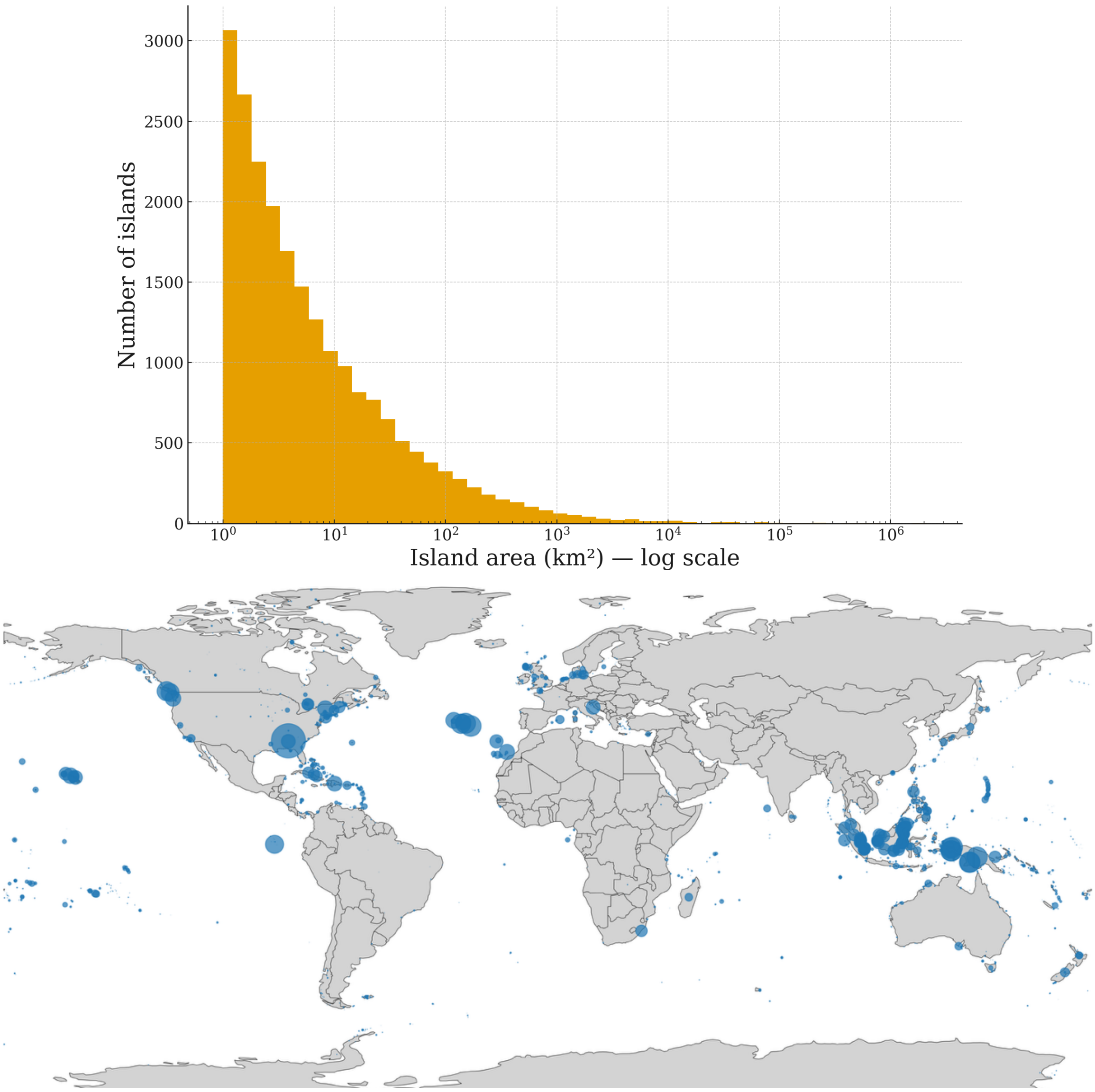


Figure S2 – Above: size distribution of all named (12,776) and unnamed Earth's islands exceeding 0.99 km$^2$. Below: sampling distribution of the dataset under study. Size of the dots represents the number of observations in a given coordinate.

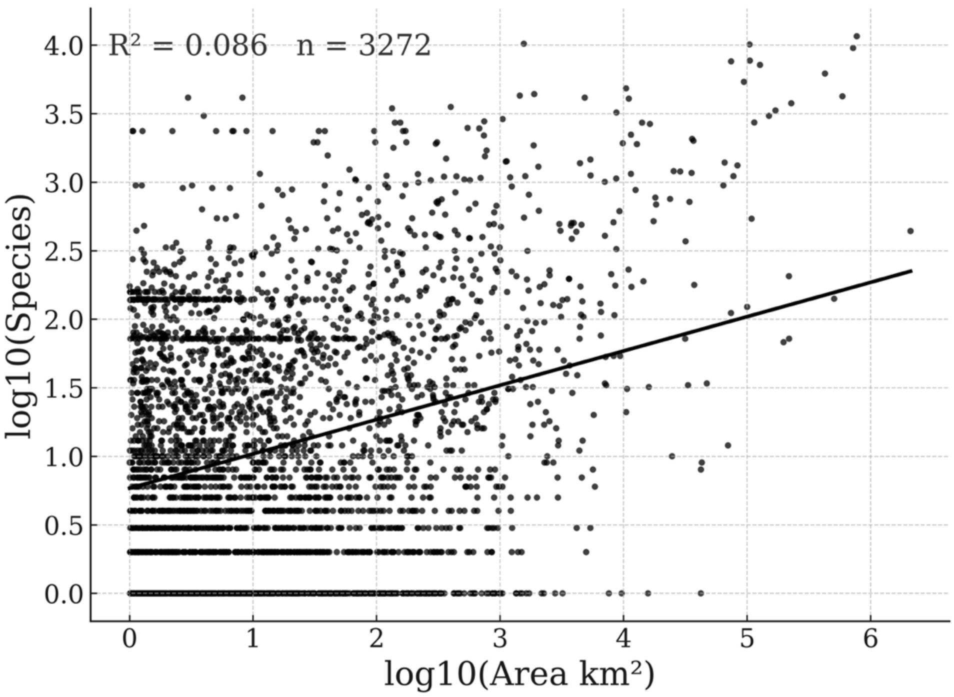


Figure S3 – Regression of island area and number of different species linked to an island in the dataset used for this study. Bigger islands retain higher biodiversities, following the IBT expectation.

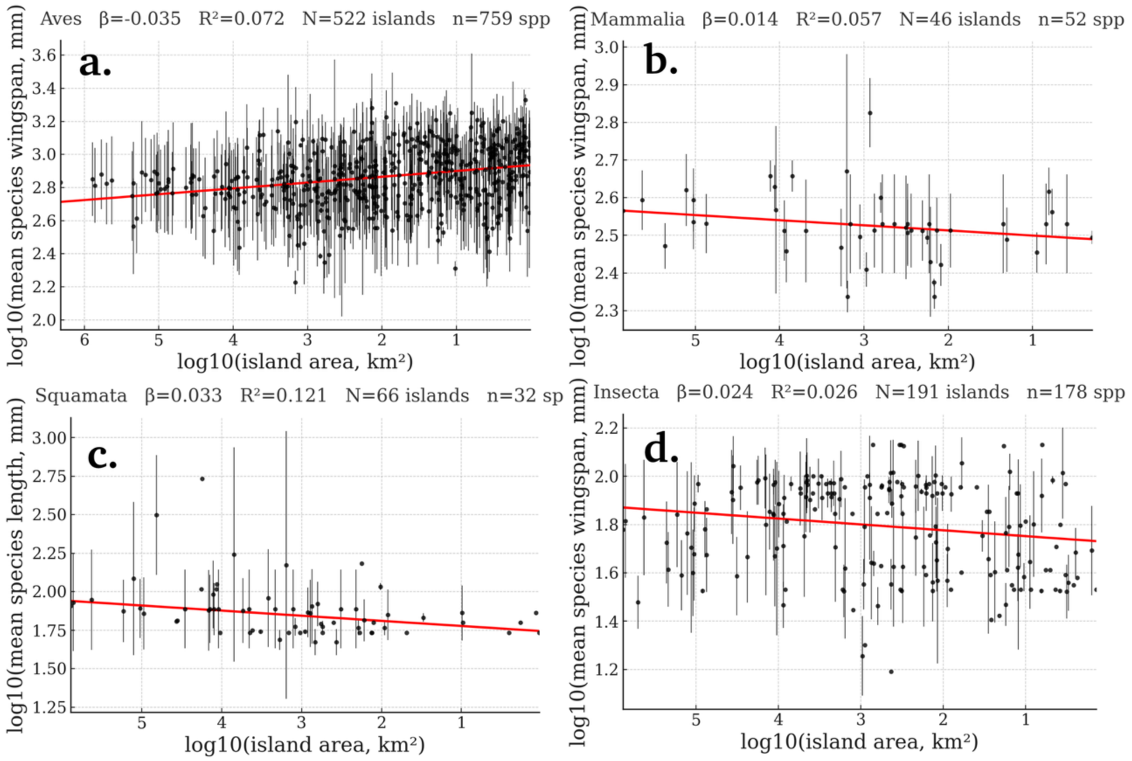


Figure S4 – Decreasing island area predicts average species length, although slope polarity is dominated by taxonomy. a: aves/birds, b: Mammals, c: scaled reptiles, d: insects. The slope is agreeing a naive IBT expectation only in a. It is worth noting that a vast majority of the length measurements in d. were carried on Coleoptera, which may suffer less isolation than other orders within insects (Peck, 2016).

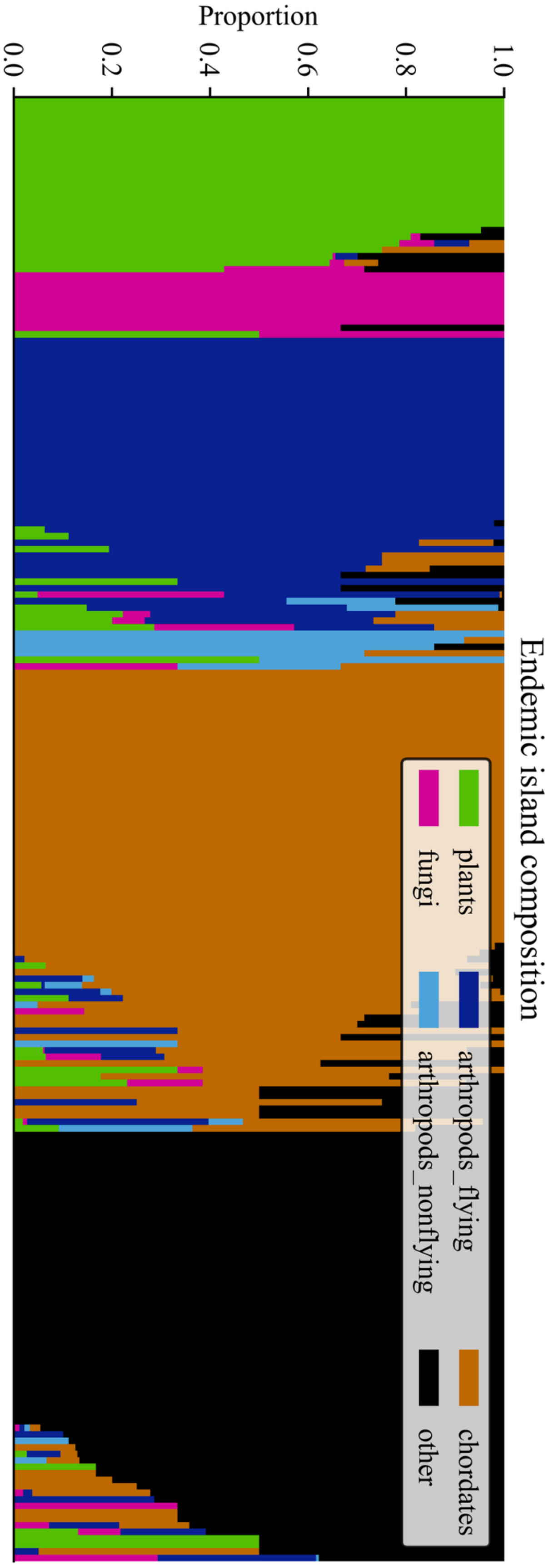


Figure S5. Genome composition in the retrieved insular dataset (only islands with endemic taxa shown). Coloured bars show 0 to 1 proportions of genome values correspondent to particular taxa. Thus, each bar represents an island. From top to bottom, we show: islands dominated by plants (green dots in main text), by fungi in pink, etc. Although the conclusions for these groups remain qualitative and subject to noise (as some island-level estimates include contributions from other taxa), the overall patterns are likely to be robust.

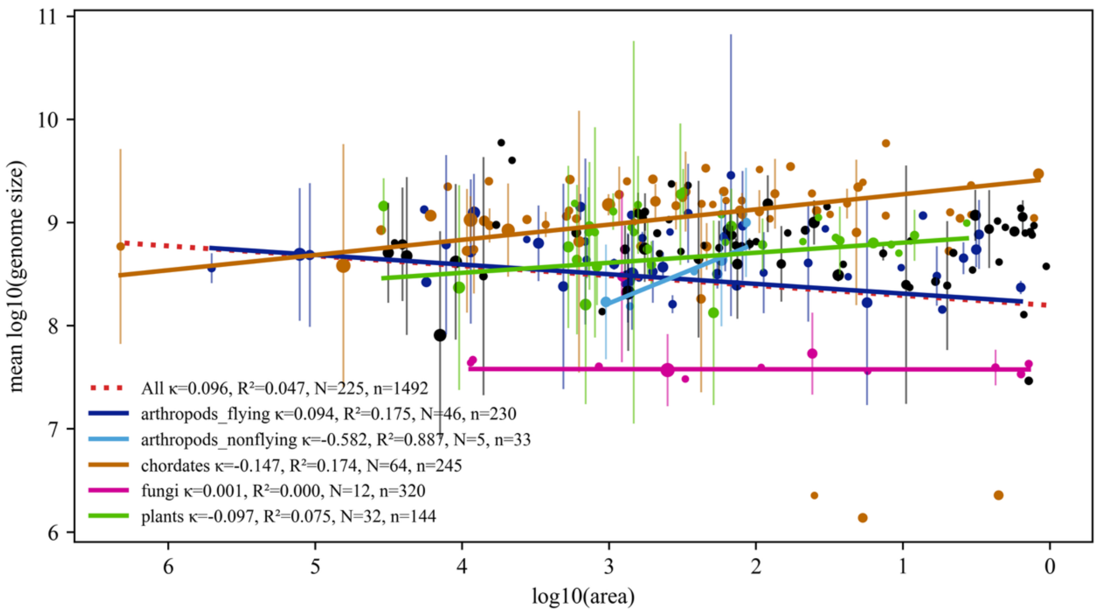


Figure S6. A reproduction of Figure 4 from main text, including standard deviations at the island level.

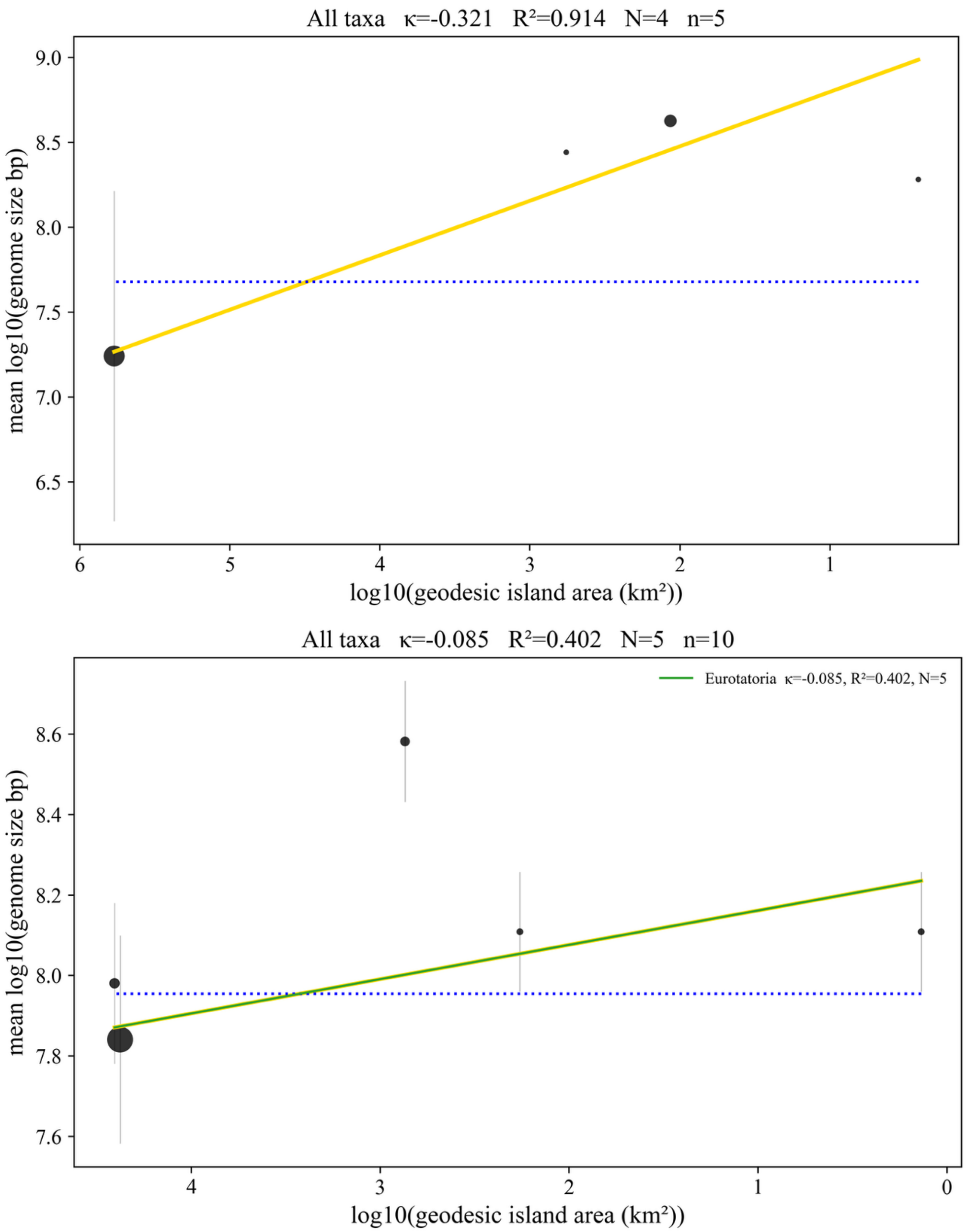


Figure S7. Endemic copepods (above) and rotifers (below) by-island level regressions show positive trends; i.e. genome size increases are predicted with decreasing island area. However, given the limited sample size, these conclusions should be regarded as preliminary.

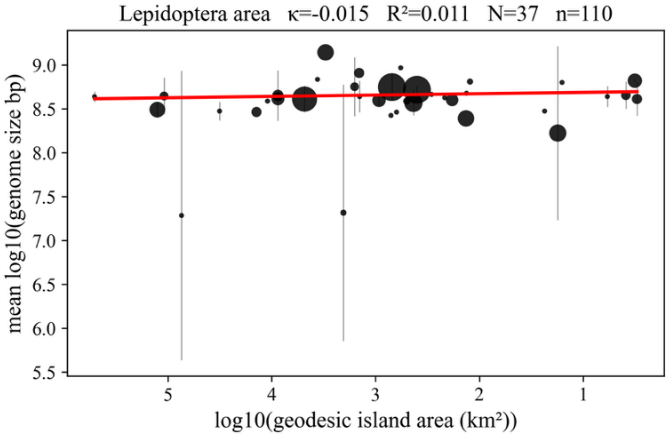


Figure S8 – Large flying insects, like beetles and moths (red regressions), show island area-dependent genomic dwarfism, with the exception of butterflies (stasis to gigantism; shown top left). Collembola (springtails; yellow regression) show genomic gigantism, as expected for non-flying arthropods.

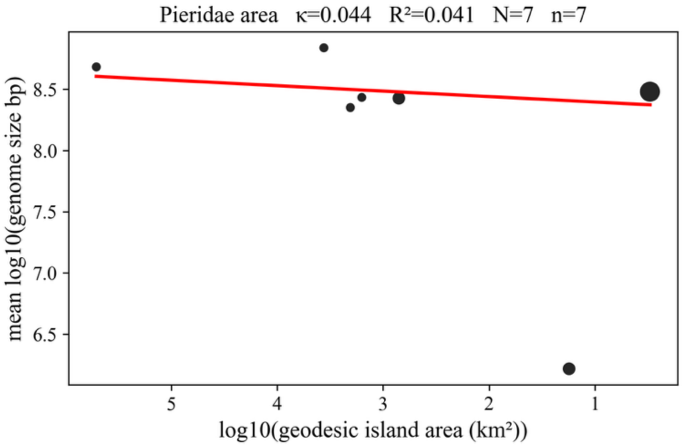


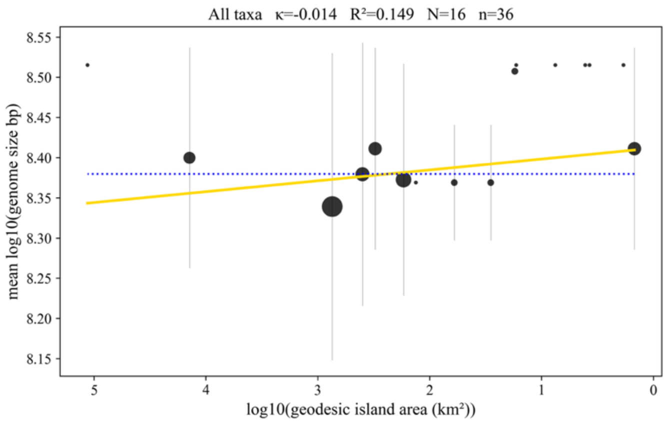


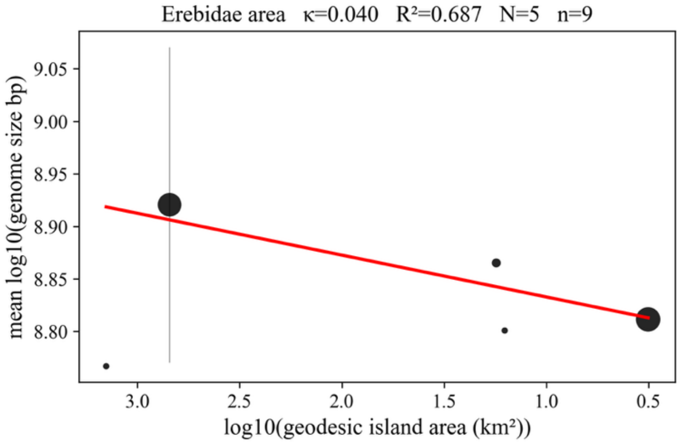


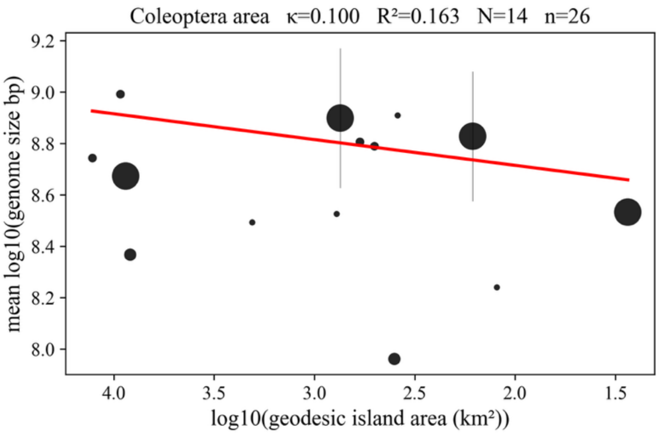

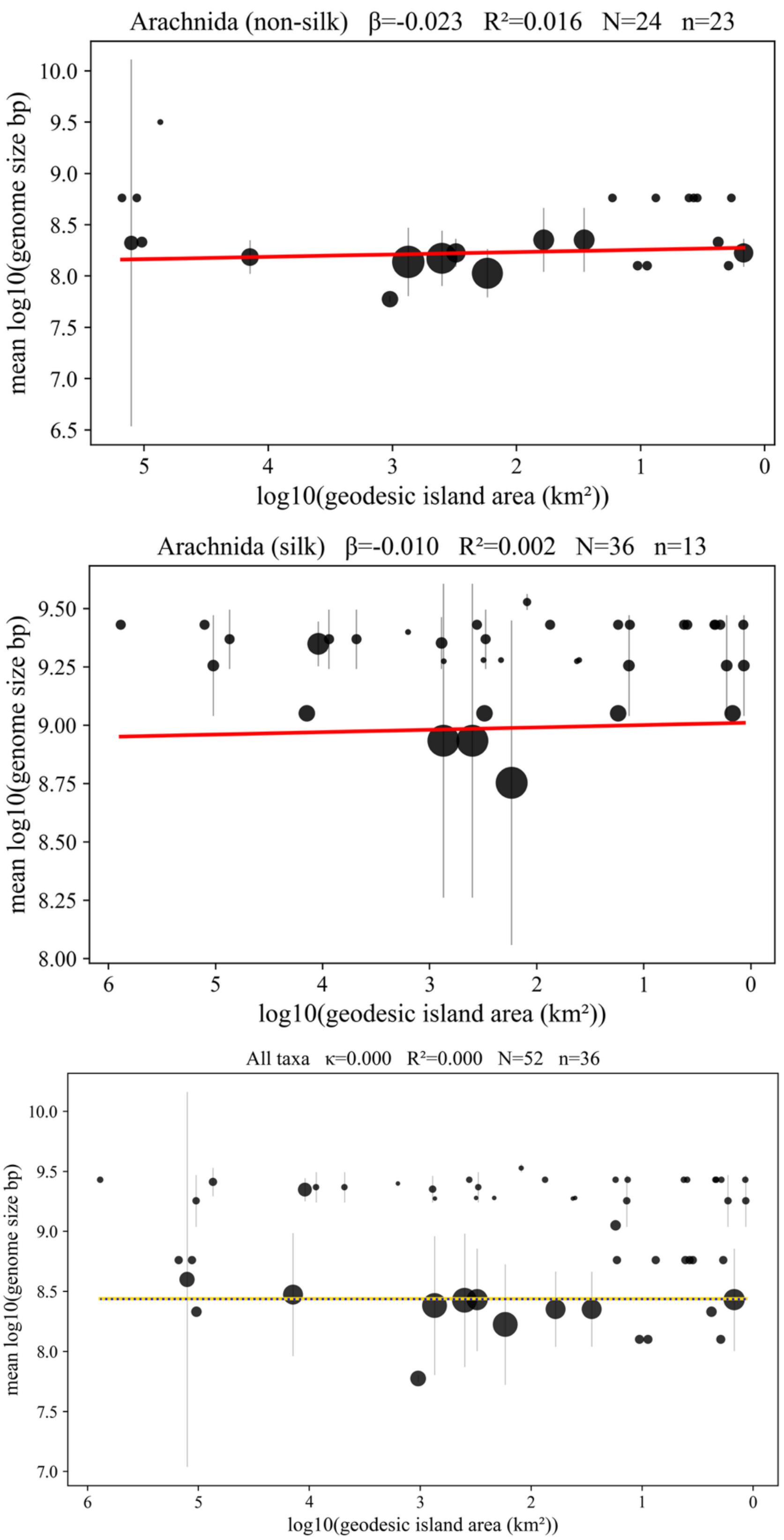


Figure S9 – Island area slightly predicts a positive change in genome size in non-ballooning species when decreasing island size (top). When analysing ballooning-capable species only, prediction is much weaker (8-fold), but in the same direction (middle). Every-one analysis is unresponsive to island area (bottom).

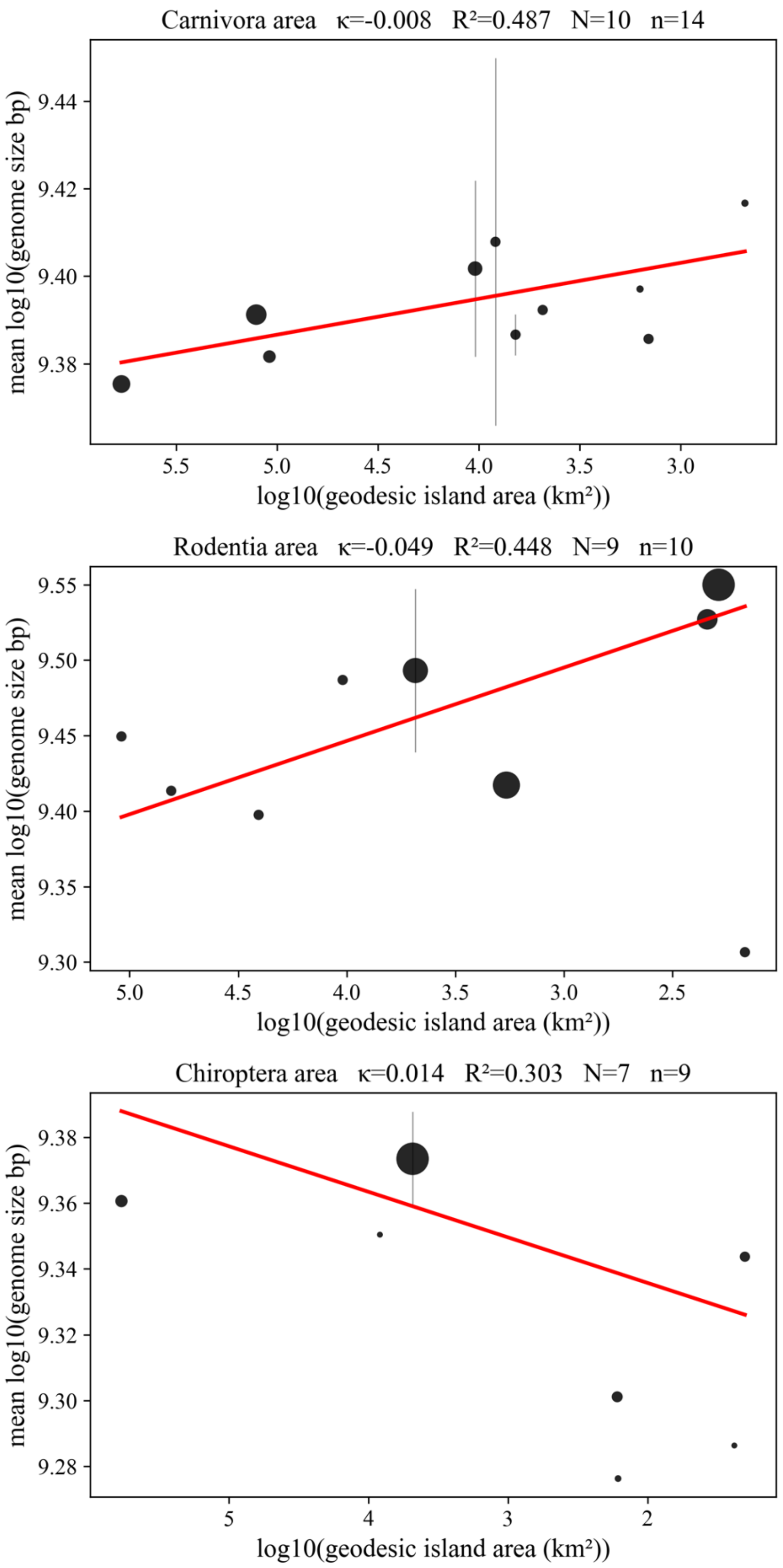


Figure S10 – Area-genome size regressions in carnivores, rodents and chiropterans.

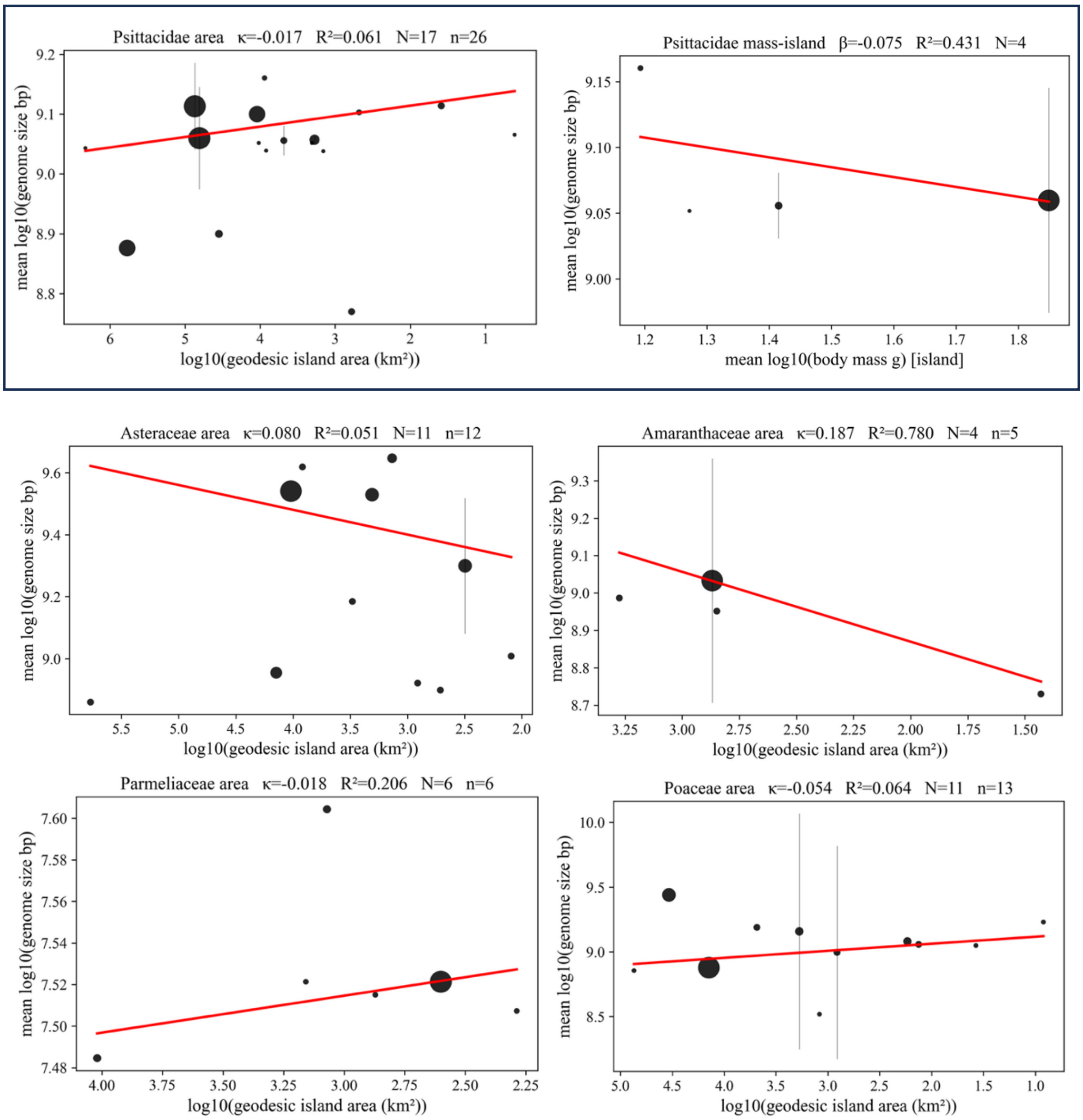


Figure S11 – Genome size-area regressions in four families of plants and one of lichens (Parmeliaceae). Top right: Psittaceae mass-area regression predicts body size dwarfism and it can be compatible with genome size gigantism if Ne is reduced.

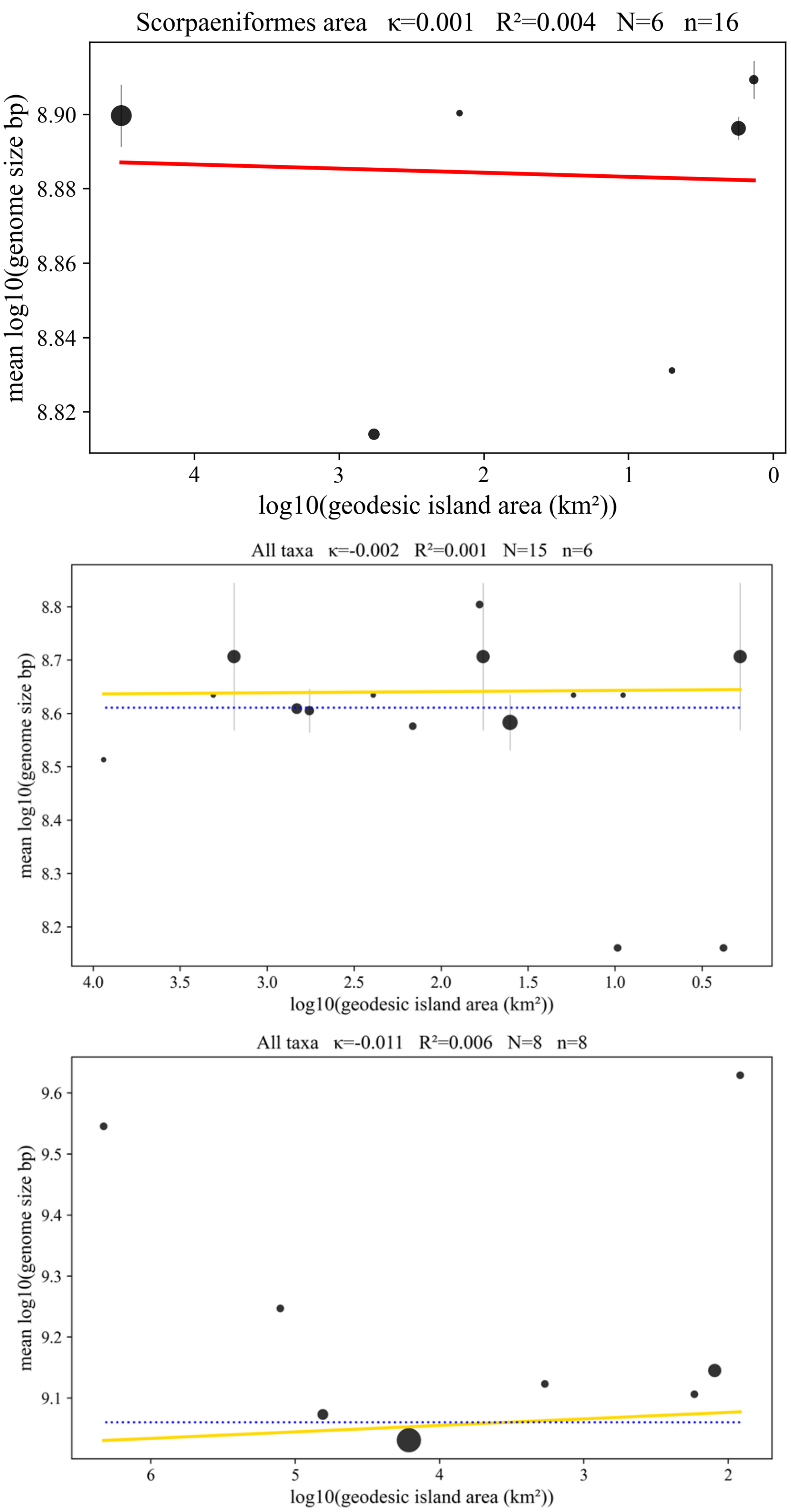


Figure S12 – Area-genome size regression in swimming, pelagic, or pedestrian species: mail-checked fishes (above), true jellyfishes (mid), malacostraca (below).

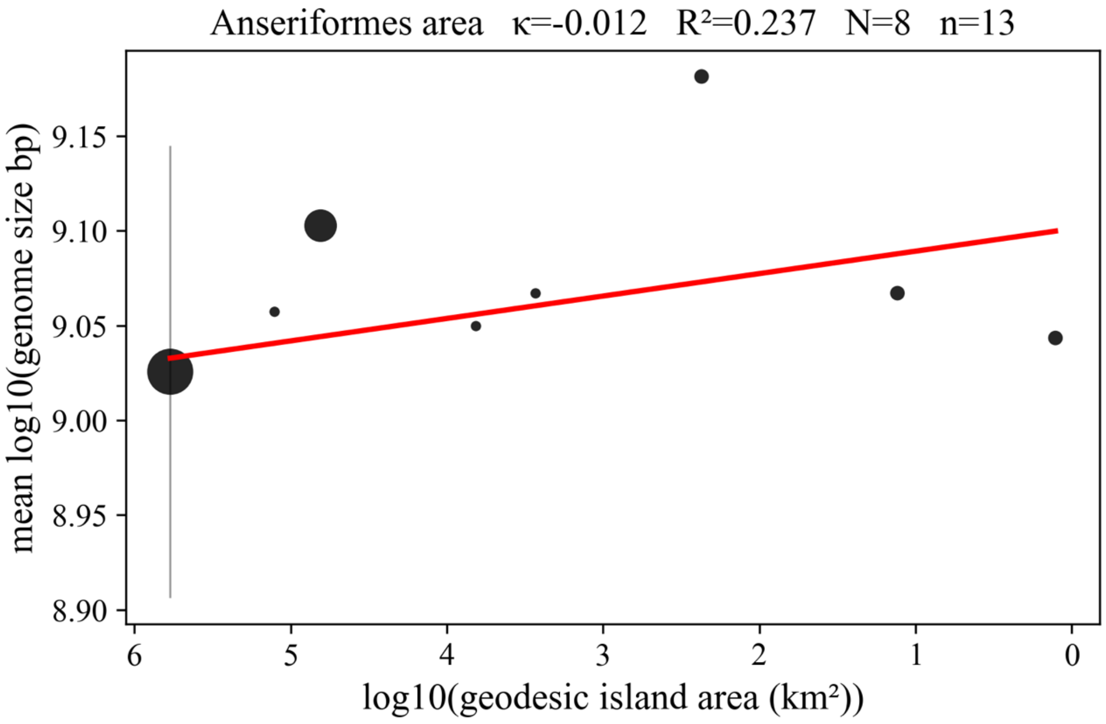


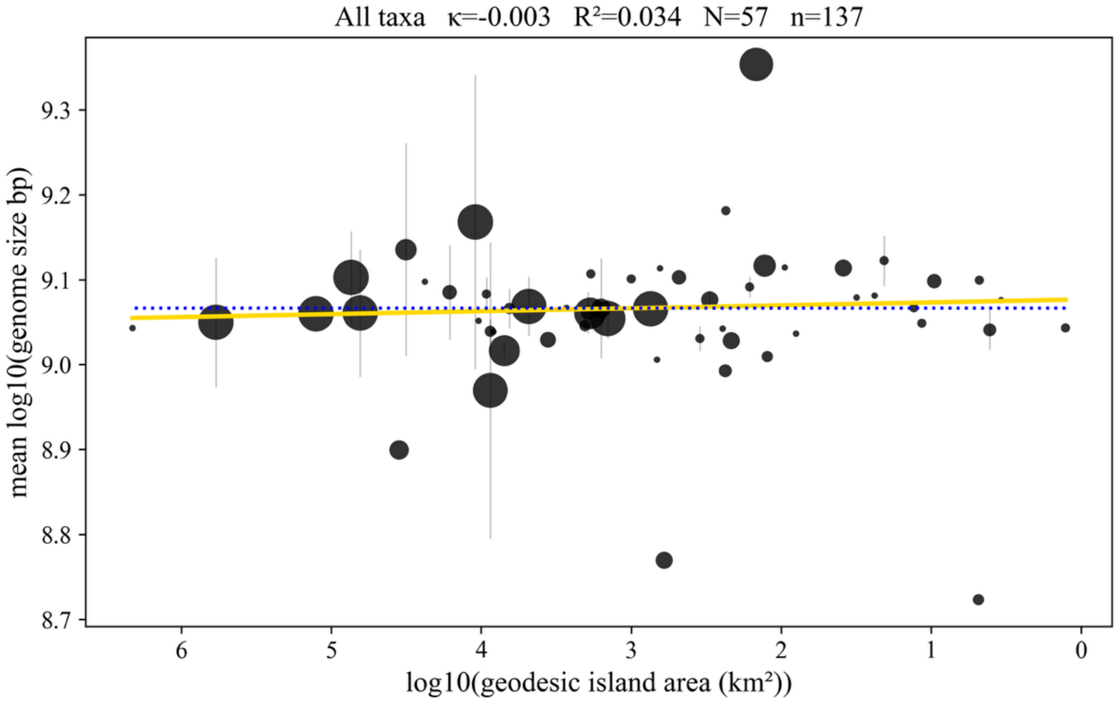


Figure S13 - Area-genome size regression in tree birds and Aves in general (below). Bigger genome sizes were predicted by the model.

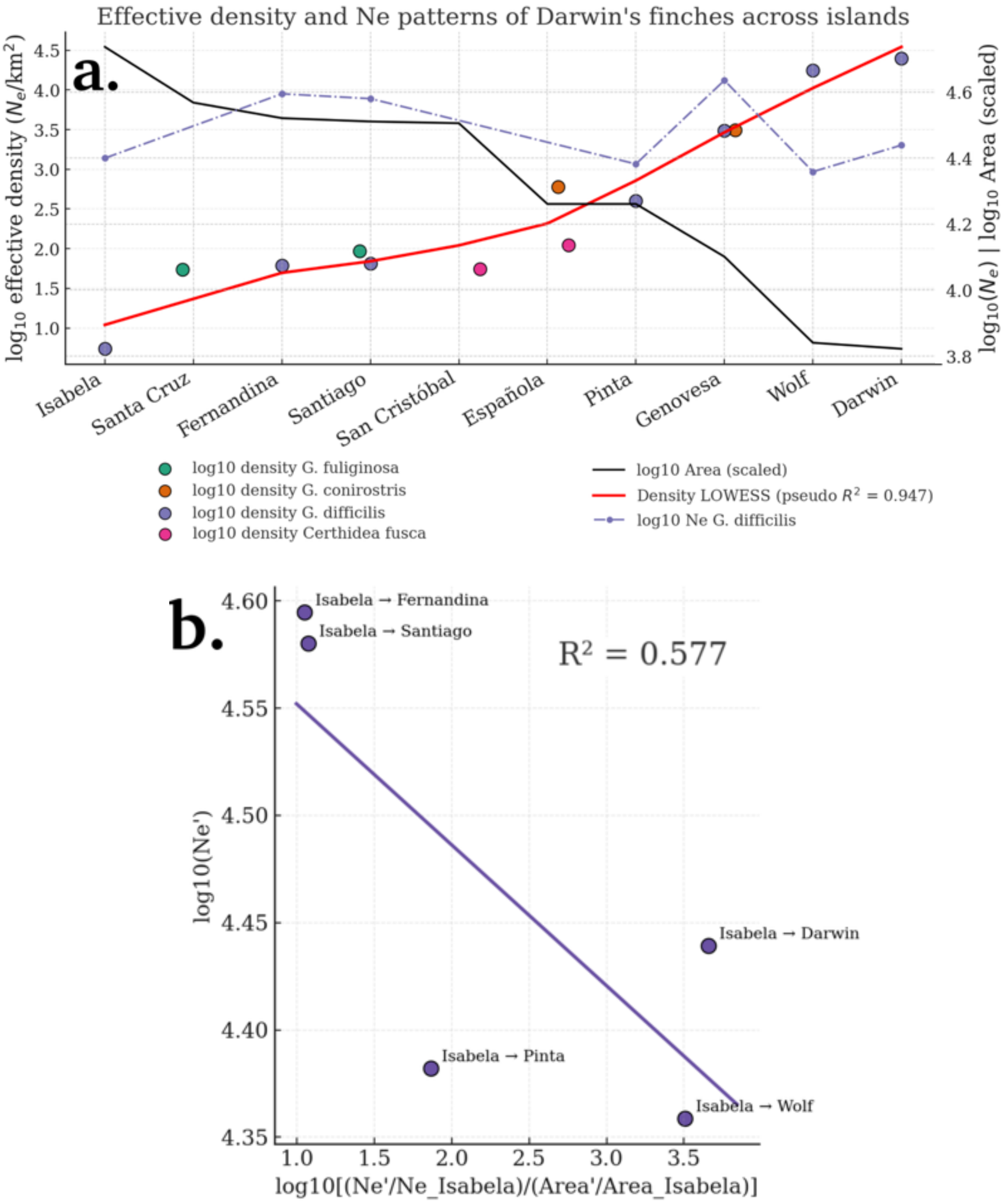


Figure S14 – Area, $N_e$ and $D_e$ of different Darwin's finches populations among the different Galapagos islands, data from (Lamichhaney et al. 2015). b. Regression of $N_e$ of G. difficile populations among the different Galapagos islands and $D_e{}'$, assuming colonisation occurred from the biggest island.

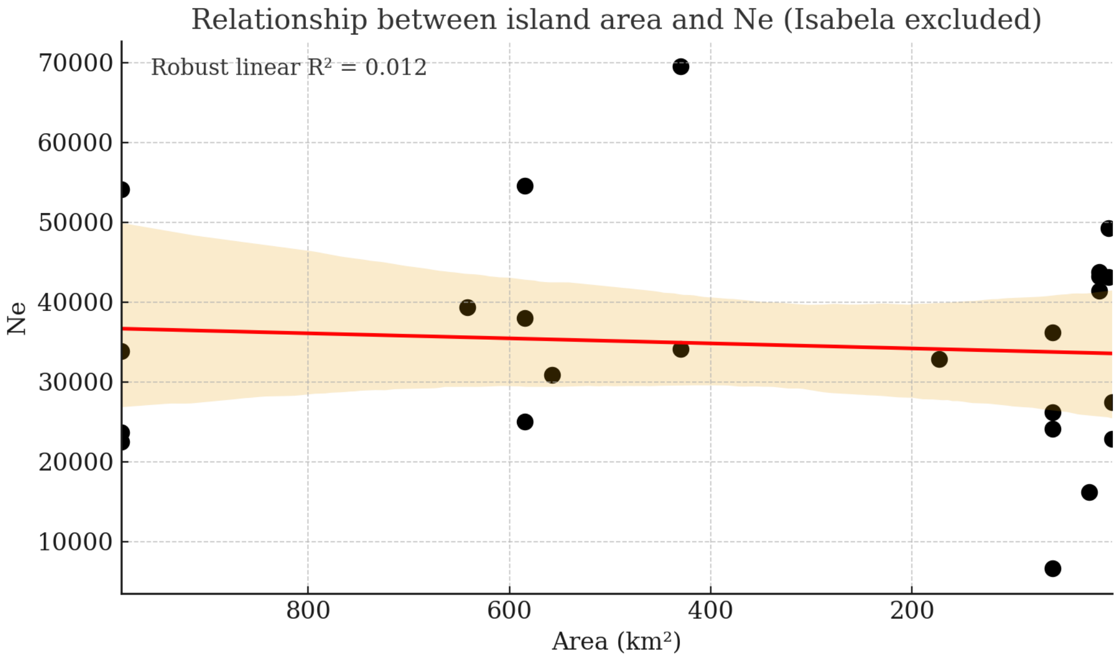


Figure S15 – Decreasing island area weakly predicts decreases in effective population size among Darwin's finches. Data from Lamichhaney et al., (2015).

Importantly, Darwin's finches should become bigger only on islands where their effective densities ($D_e$') decrease, if $\Delta N_e > \Delta$area. Conversely, they should become smaller where colonisation of a new island results in $\Delta N_e < \Delta$area. Although we lack body-size information for these particular finch populations and therefore cannot test whether this expectation is fulfilled, the more general result suggested island area-dependent gigantism in birds (Fig. 2a in main). If $\varkappa > 0$ is fulfilled for Darwin's finches with increasing $D_e$', the model cannot be correct unless significant gene flow occurs among islands. In that case, island area would be a poor predictor of the area effectively occupied by these finches.

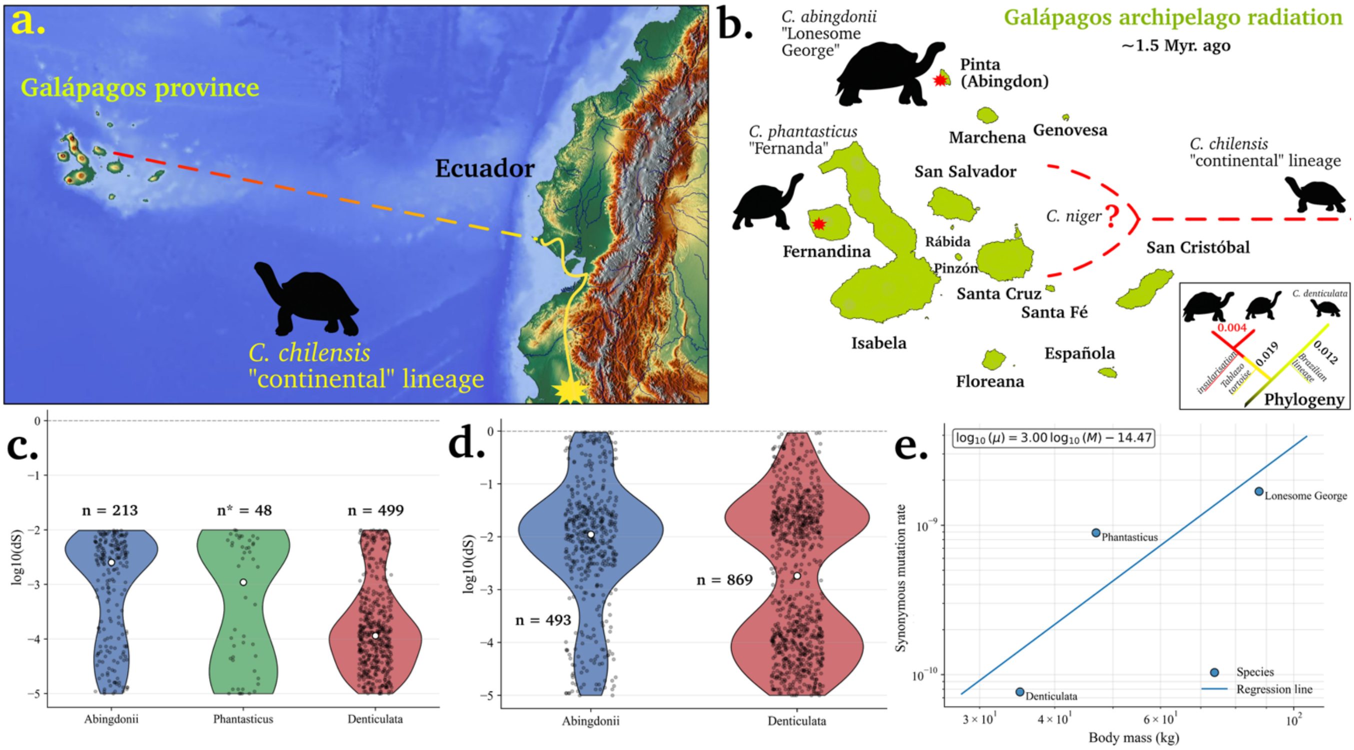


Figure S16. a. Chelonoidis lineage radiation from South America to the Galápagos archipelago. b. Selection of three genomes from the Chelonoidis phylogeny for the approximation of branch-specific synonymous mutation rates, based on effective area and body size. The closest available reference genome from an extant continental population was *C. denticulata*. The insular genomes belonged to a *C. phantasticus* individual, "Fernanda", and the last known individual of the putatively extinct *C. abingdonii*, "Lonesome George". The three species differ in body weight and inhabit landmasses of different sizes. c. The lower ranges of the dS distributions among shared orthologues ($10^{-5} \leq$ dS $\leq 10^{-2}$) agree with the model. n*: only 48 orthogroups were considered reliable in the *C. phantasticus* assembly, which had approximately 5x coverage. d. Full-range dS distributions for the insular and continental lineages (Wilcoxon signed-rank test: $p = 2.05 \times 10^{-4}$; paired t-test: $p = 7.43 \times 10^{-4}$). e. The regression between body mass and lower-bound mutation rates follows a power law consistent with the model: when reduced area lowers effective density, increased body size predicts higher mutational loads.

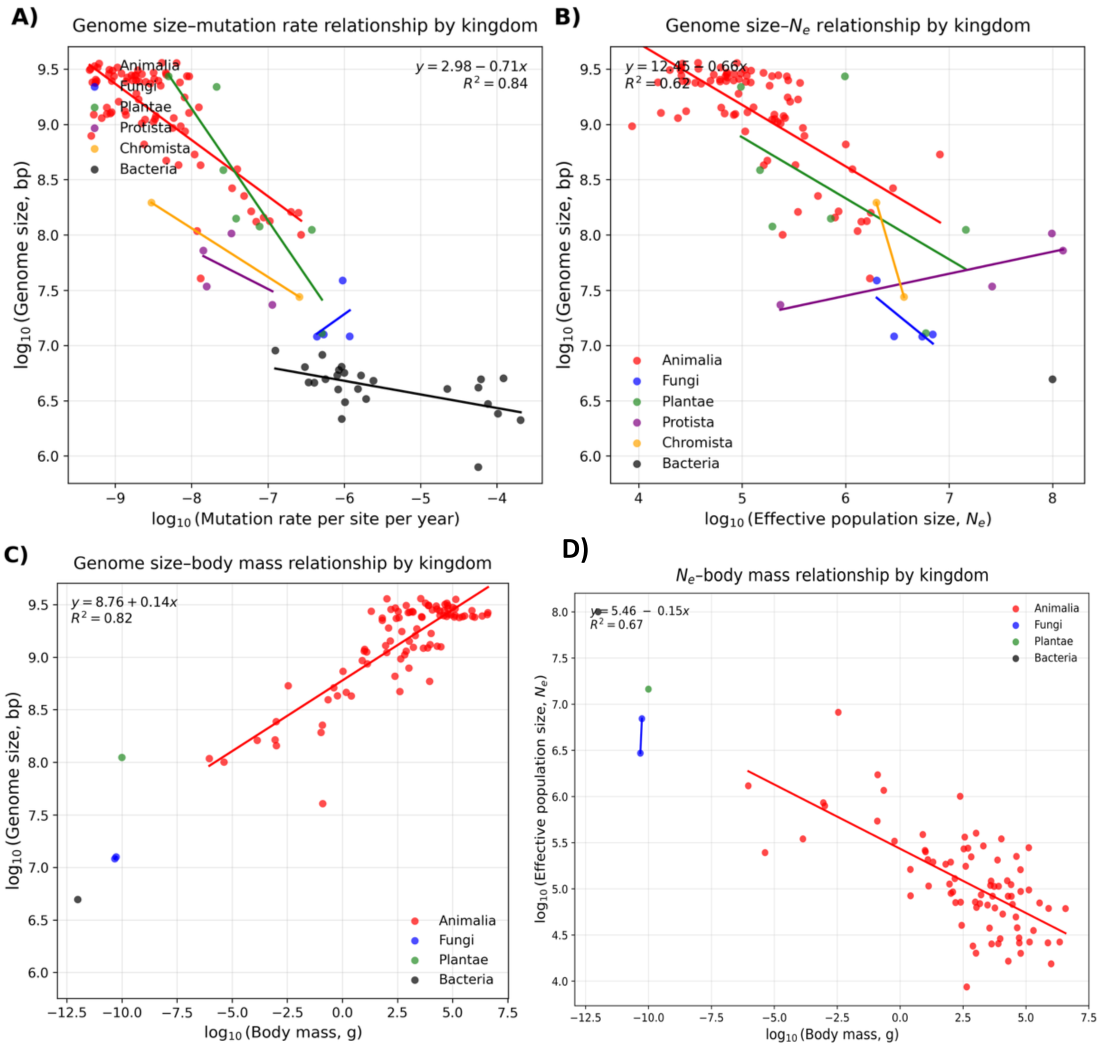


Figure S17. Regression between genome size and mutation rate, effective population size, and body mass from a global dataset. As shown, Ne and body mass allow for prediction of genome size.
